# The FXR1 network acts as signaling scaffold for actomyosin remodeling

**DOI:** 10.1101/2023.11.05.565677

**Authors:** Xiuzhen Chen, Mervin M. Fansler, Urška Janjoš, Jernej Ule, Christine Mayr

## Abstract

It is currently not known whether mRNAs fulfill structural roles in the cytoplasm. Here, we report the FXR1 network, an mRNA-protein (mRNP) network present throughout the cytoplasm, formed by FXR1-mediated packaging of exceptionally long mRNAs. These mRNAs serve as underlying condensate scaffold and concentrate FXR1 molecules. The FXR1 network contains multiple protein binding sites and functions as a signaling scaffold for interacting proteins. We show that it is necessary for RhoA signaling-induced actomyosin reorganization to provide spatial proximity between kinases and their substrates. Point mutations in FXR1, found in its homolog FMR1, where they cause Fragile X syndrome, disrupt the network. FXR1 network disruption prevents actomyosin remodeling—an essential and ubiquitous process for the regulation of cell shape, migration, and synaptic function. These findings uncover a structural role for cytoplasmic mRNA and show how the FXR1 RNA-binding protein as part of the FXR1 network acts as organizer of signaling reactions.

## Introduction

Cells use biomolecular condensates to generate compartments that are not surrounded by membranes^1^. These compartments are thought to enable the spatial organization of biochemical activities^2^. For example, condensates function as signaling clusters for T cell activation or concentrate factors for the nucleation and assembly of actin filaments^3,4^. Cytoplasmic messenger ribonucleoprotein (mRNP) granules are a group of condensates, formed through self-assembly of mRNAs and their bound proteins. They include P bodies and stress granules and are thought to function in mRNA storage and decay^5,6^, where it appears that mRNAs take on passive roles of being stored or degraded. In contrast, within TIS granules, mRNAs actively contribute to protein functions by establishing mRNA-dependent protein complexes^7–9^. An apparent difference between P bodies or stress granules and TIS granules is the network-like morphology of TIS granules, which is generated through RNA-RNA interactions^7,10^. In this study, our goal was to identify another cytoplasmic mRNP network and to investigate whether mRNAs have broader structural or regulatory roles in addition to serving as templates for protein synthesis.

We focused our study on FXR1, an RNA-binding protein from the family of Fragile X-related (FXR) proteins^11^. FXR proteins are ancient and were found in invertebrates but have expanded into three family members in vertebrates^11,12^. *FXR1* and *FXR2* are homologs of the *FMR1* gene, whose loss of function causes the most common form of hereditary mental retardation in humans, Fragile X syndrome (FXS)^11,13^. FXR1 has recently also been implicated in neurological disorders, as several genome-wide association studies found variants in *FXR1* that are associated with a higher risk for autism spectrum disorder (ASD), intellectual disability, and schizophrenia^14–17^.

FXR1 is an essential gene in humans, as loss of function of FXR1 is not tolerated^18^. Whereas mice with knockouts of *FMR1* or *FXR2* are viable, loss of FXR1 results in perinatal lethality, likely due to cardiac or respiratory failure^19^. FXR1 has mostly been studied as a regulator of translation in brain, testis, and muscle^20–22^. However, *FXR1* may have broader roles, as *FXR1* is ubiquitously expressed and was detected among the top 15% of expressed genes in primary fetal and adult cell types (Fig. S1A)^23^.

Here, we find that the longest expressed mRNAs assemble with FXR1 into a large cytoplasmic mRNP network, which we call the FXR1 network. Only a small fraction of FXR1 is stably bound to mRNA, these FXR1 molecules together with the bound mRNAs act as network scaffold. FXR1 contains multiple protein binding sites, including coiled-coil (CC), Tudor, and RGG domains, which allow the recruitment of most FXR1 molecules as clients into the network, thus generating a high FXR1 concentration. Additional clients, such as signaling molecules with similar protein domains as FXR1 are also recruited to the network, which promotes their proximity. We show that an intact FXR1 network is necessary for RhoA signaling-induced actomyosin reorganization, as it provides proximity between the Rho-associated kinase and its substrates. Actomyosin remodeling is crucial for many cellular processes including the control of cell shape, migration, and synaptic function. Taken together, we demonstrate that mRNAs fulfil structural roles in the cytoplasm. They provide an underlying scaffold for FXR1, whose high concentration of multiple protein binding sites generates a platform for signaling molecules to utilize this mRNP network despite lacking RNA-binding domains.

## Results

### FXR1 and its bound mRNAs assemble into a large cytoplasmic mRNP network

To identify additional cytoplasmic mRNP networks, we performed a small-scale high-resolution imaging screen on highly abundant cytoplasmic RNA-binding proteins. Using immunostaining, we observed that endogenous FXR1 forms a network-like structure that covers the whole cytoplasm (Fig. 1A). We call this assembly the FXR1 network, which is composed of extensively connected spherical granules (Fig. 1A). The FXR1 network is present in all cells of all eight cell types examined (Fig. S1B) and was also observed in C2C12 myotubes^24^.

**Figure 1.**
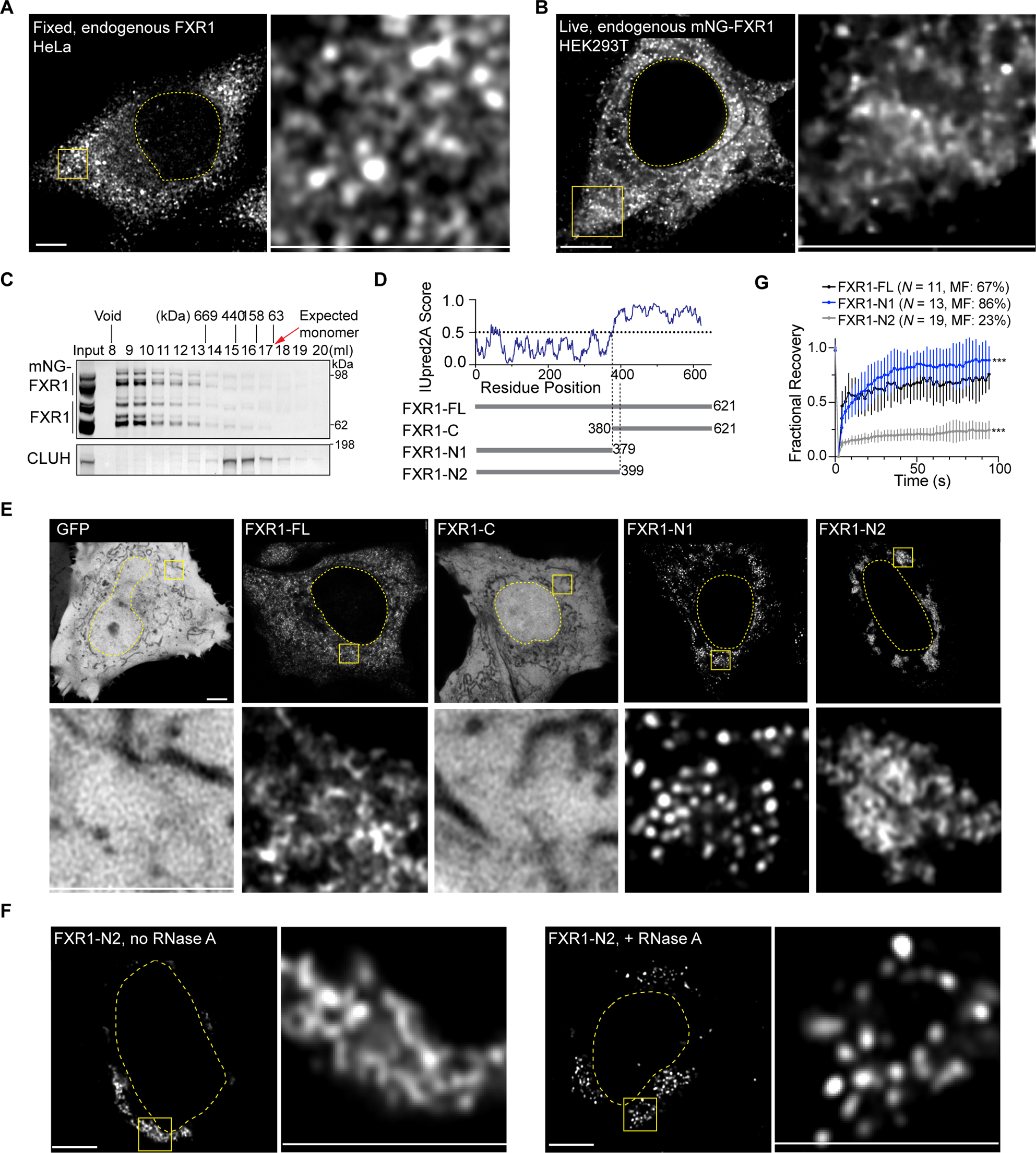
FXR1 assembles with its bound mRNAs into a cytoplasmic mRNP network. **A.** Immunofluorescence staining of endogenous FXR1 protein in HeLa cells. The dotted line indicates the nucleus. Right panel is a zoomed-in image of the region in the yellow box. All cells contain the network, and a representative confocal image is shown. All scale bars in Figure 1 are 5 µm. **B.** Live cell confocal imaging of HEK293T cells with endogenous monomeric NG-tagged FXR1 protein. All cells contain the network and a representative image is shown. **C.** Size exclusion chromatography of cells from (**B**), immunoblotted for FXR1. CLUH was used as loading control. mNG-FXR1 and FXR1 have the same elution pattern. **D.** IUpred2A score of human FXR1. A score greater than 0.5 indicates an IDR. Schematics of GFP-fusion constructs. The numbers denote amino acids. **E.** Live cell confocal imaging of HeLa cells transfected with the FXR1 constructs from (**D**) after knockdown of endogenous FXR1. The GFP fIuorescence pattern shown for each construct was observed in all cells expressing the respective FXR1 constructs. Representative images are shown. See Fig. S4C for quantification. **F.** Confocal imaging of HeLa cells transfected with GFP-FXR1-N2 after digitonin permeabilization in the presence or absence of RNase A treatment for 30 minutes. Representative images from at least three independent experiments are shown, where 21 cells were examined. **G.** FRAP analysis of GFP-FXR1-FL, -N1, and -N2 expressed in HeLa cells. Shown is a normalized FRAP curve as mean ± std from at least 11 cells each. MF, mobile fraction. See Videos S7-9 for representative fluorescence recovery. Mann-Whitney test, N1 vs. FL, ***, *P<*10^−21^; N2 vs. FL, ***, *P<*10^−165^.

The network-like morphology was also observed with live cell imaging of monomeric NeonGreen (NG)-tagged endogenous FXR1 (Fig. 1B, S1C-G). Both major splice isoforms expressed in non-muscle cells are capable of FXR1 network formation (Fig. S2A-F). The higher-order assembly of FXR1 observed by imaging was confirmed using size-exclusion chromatography (Fig. 1C). FXR1 protein exists predominantly within high-molecular-weight complexes with an estimated size of more than 1,000 kDa. In contrast, monomeric FXR1 is present at very low levels in cells (Fig. 1C).

### The underlying scaffold of the FXR1 network is RNA

To learn how FXR1 assembles into a network, we ectopically expressed monomeric GFP-fused FXR1 and its variants in cells depleted of endogenous FXR1 protein (Fig. S2E). In its N-terminal half, FXR1 protein contains several folded domains that are followed by an intrinsically disordered region (IDR) (Fig. 1D). Expression of the IDR fused to GFP resulted in diffusive signal, similar to that of GFP alone, whereas expression of the folded domains, which contain two KH RNA-binding domains^11^, generated spherical granules, different from the full-length FXR1 protein (Fig. 1E). KH domain mutation generated a diffusive signal, indicating that formation of the granules requires RNA binding of FXR1 (Fig. S3A-C).

Intriguingly, when fusing the first 20 aa of the IDR with the folded domains of FXR1, the spherical granules turned into a network-like structure (FXR1-N2, Fig. 1E). The first 20 aa of the IDR contain an RG-rich region. RG motifs are known as RNA-binding regions^25^, suggesting that RNA may be responsible for connecting the granules and for network formation (Fig. S3C). We tested this prediction by treating the assembled network with RNase A, which reverted the network into spherical granules (Fig. 1F, S3D). Furthermore, mutating the five arginines of the RG motif into alanines abolished its ability to connect the granules. In contrast, substituting the arginines with five positively charged lysine residues retained this activity (Fig. S3C). These results indicate that RNA forms the connections between the spherical granules. Together, these data demonstrate that RNA is essential for both the initial granule formation and their remodeling into a network.

### The FXR1 network is highly dynamic

To better understand the material properties of the FXR1 network, we acquired time lapses and performed fluorescent recovery after photobleaching (FRAP). Full-length FXR1 generates a highly dynamic network, whose components are highly mobile, as over 50% of the initial fluorescence recovered in less than two seconds (Fig. 1G, Videos S1 and S2). In contrast, FXR1-N2 localizes to the perinuclear region, where it forms a rather static assembly, with a low fraction of mobile molecules, according to FRAP (Fig. 1G, Video S3). These results suggest that the IDR is responsible for the high protein mobility in the network. FXR1-N1 generates highly dynamic and mobile granules that rarely fuse upon contact (Figure 1G, Videos S4 and S5). Over twelve hours, their numbers and occupied areas remain quite constant (Fig. S3E, S3F). In the presence of the RG motif however (FXR1-N2), the granule numbers decrease substantially, while their sizes increase (Fig. S3G, S3H), indicating that the RG motif is responsible for the fusion of the granules and network formation.

### FXR1 dimerization through the CC domains nucleates the FXR1 network

Although FXR1 is primarily known as an RNA-binding protein, it also contains multiple domains for protein:protein interactions (Fig. 2A)^11,26,27^. FXR1 contains two Tudor domains, which mediate dimerization and bind to methylated arginines^25,28–30^. The two KH RNA-binding domains have also been reported as protein:protein interaction domains^11,31–34^. The KH0 domain may act as protein:protein interaction domain because it lacks the GXXG motif required for RNA-binding^27^. FXR1 contains two predicted CC domains (Fig. S4A)^26,35^. Within its IDR, there are three arginine-rich regions (RG, RGG, R). RG/RGG motifs are multifunctional as they can bind to RNA or to protein^25,36–38^. They often bind to other RGG motifs, resulting in homo- or heterooligomerization^25,28,29,37–39^. Taken together, FXR1 contains at least five domains for protein:RNA interactions and nine domains capable of forming protein:protein interactions (Fig. 2A).

**Figure 2.**
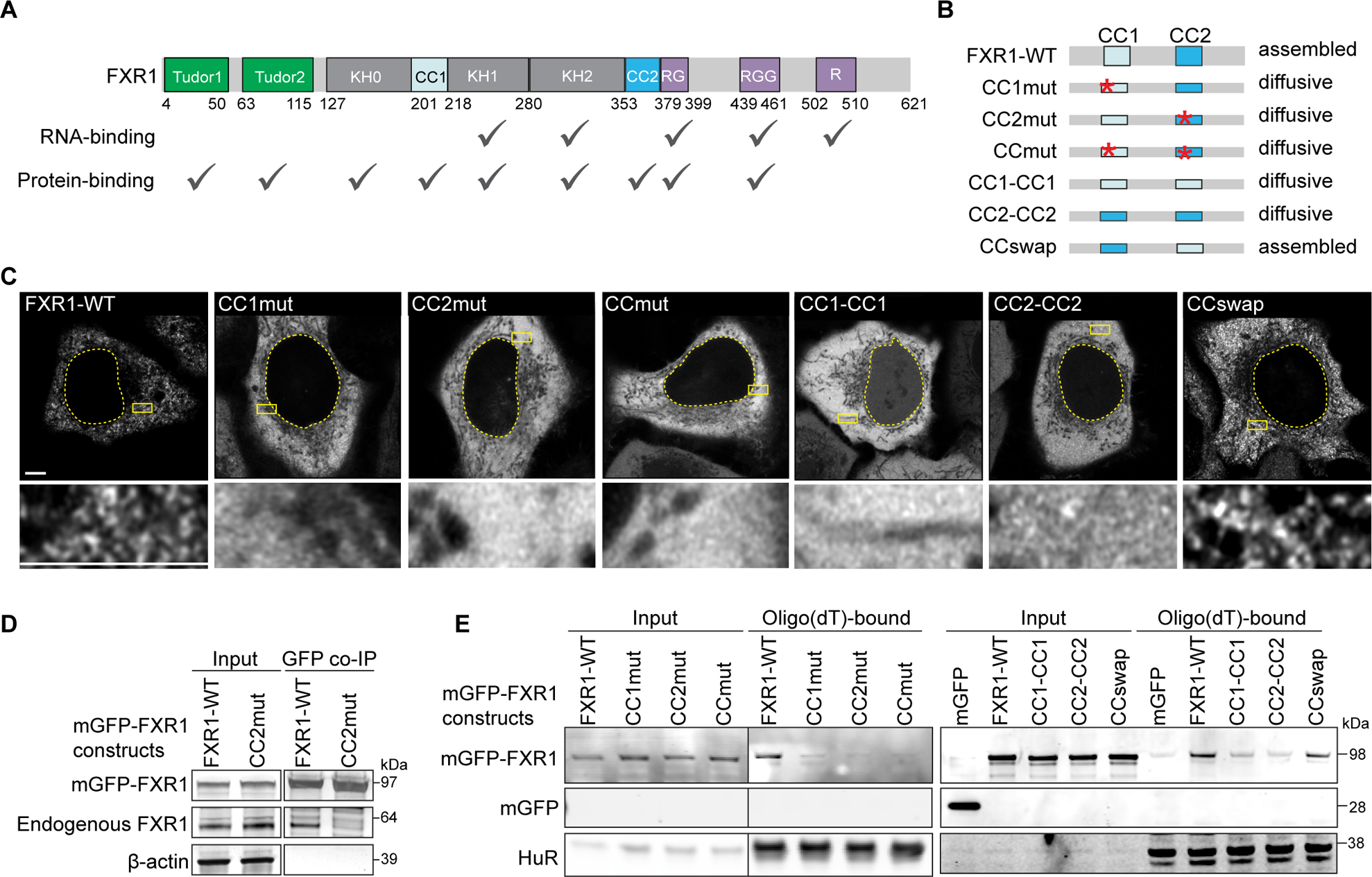
FXR1 dimerization through the CC domains promotes mRNA binding and nucleates the FXR1 network. **A.** Amino acid boundaries of FXR1 protein domains. Domains capable of binding to RNA or protein are indicated. **B.** Schematic of FXR1 CC mutant constructs and their resulting FXR1 network assembly states. Red star symbols represent single point mutations. CC1mut is N202P, CC2mut is V361P. See Fig. S4A-E for details. **C.** Live cell confocal imaging of HeLa cells transfected with GFP-FXR1 constructs from (**B**) after knockdown of endogenous FXR1, shown as in Fig. 1A. Representative images from at least three independent experiments are shown where 38 cells were examined. See Fig. S4C for quantification. Scale bar, 5 µm. **D.** GFP co-IP of endogenous FXR1 after ectopic expression of GFP-FXR1-WT or GFP-FXR1-CC2mut in HeLa cells. Actin is shown as loading control. 1% of input was loaded. **E.** Oligo(dT) pulldown, performed without cross-linking, of mRNA-bound FXR1 in FXR1/FXR2/FMR1 triple KO U2OS cells after ectopic expression of GFP or GFP-FXR1 constructs from (**B**). The endogenously expressed RNA-binding protein HuR was used as positive and loading control for oligo(dT)-bound proteins. 2.5% and 5% of input were loaded in the left and right panels, respectively.

To probe the molecular mechanism of FXR1 network assembly, we set out to generate an FXR1 assembly-deficient mutant, while keeping the RNA-binding domains intact. Removal of the Tudor domains resulted in network disruption that was restored upon overexpression (Fig. S3I), suggesting that the Tudor domains are not essential for network assembly. In contrast, intact CC domains are essential to nucleate the FXR1 network (Fig. 2B, 2C). Introduction of a single helix-breaking point mutation in either one of the CC domains was sufficient to fully disrupt the FXR1 network (Fig. 2B, 2C, S4A-D)^40^. Moreover, FXR1 variants that contained only either CC1 or CC2 at both positions could not nucleate the FXR1 network, whereas swapping the CC domains maintained FXR1 network assembly (Fig. 2B, 2C, S4C). These results converge on a model wherein FXR1 network formation requires heteromeric binding of the two CC domains, which is supported by biochemical evidence that intact CC domains are essential for dimerization of FXR1 (Fig. 2D).

### FXR1 dimerization strongly promotes mRNA binding

Since RNA binding is essential for network assembly (Fig. 1F, Fig. S3A-D), we determined whether FXR1 dimerization affects its mRNA binding capacity. We performed native oligo(dT) pulldown experiments using GFP-tagged wildtype (WT) FXR1 or the CC mutants expressed at levels similar to the endogenous protein (Fig. S2E)^24,41^. Only WT FXR1 stably interacted with mRNA (Fig. 2E). In contrast, mRNA binding of the FXR1 CC mutants was strongly reduced, indicating that monomeric FXR1 is a poor mRNA-binding protein. As swapping the CC domains rescued RNA binding, these data indicate that FXR1 dimerization is required for stable mRNA binding in cells (Fig. 2E). When comparing mRNA binding of FXR1 with that of HuR, we observed that nearly all of HuR was enriched by oligo(dT) pulldown, but only a small fraction of FXR1, estimated to be ∼2%, was bound to mRNA (Fig. 2E). mRNA binding to FXR1 enables higher-order assembly of FXR1, as indicated by size exclusion chromatography, which showed that FXR1 with mutated CC domains is predominantly present as monomeric protein in cells (Fig. S4E). These data indicate that FXR1 dimerization is a prerequisite for stable RNA binding. They also suggest that only a minority of FXR1 is stably bound to RNA, whereas most FXR1 molecules associate with the network in an RNA-independent manner.

### FMR1 also forms a large cytoplasmic mRNP network

The FXR family member FMR1 has the same domain architecture as FXR1 (Fig. 3A). Endogenous FMR1 is also present as a network in cells (Fig. S4F, S4G). In HeLa cells, the FXR1 and FMR1 networks partially overlap (Fig. S4G-I). Similar to FXR1, the folded domains in the N-terminus were sufficient for formation of spherical granules and addition of the RG domain of the IDR connected the granules and induced network formation (Fig. S4J, S4K). FMR1 also required intact CC domains for network assembly and stable RNA binding (Fig. 3B, 3C).

**Figure 3.**
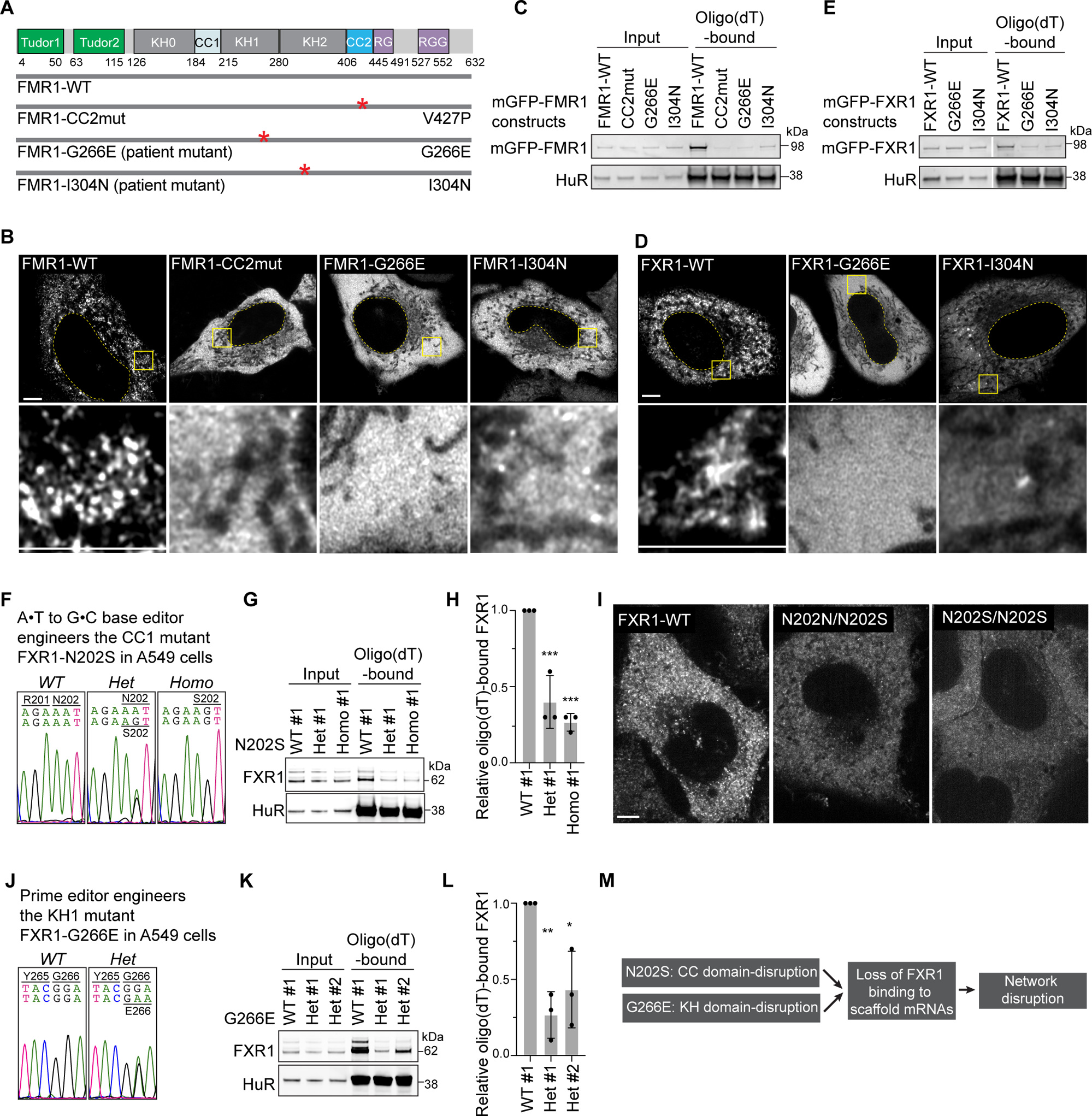
The FXS mutations I304N and G266E disrupt the FMR1 and FXR1 networks. **A.** Amino acid boundaries of FMR1 protein domains and schematics of FMR1 constructs. **B.** Live cell confocal imaging of HeLa cells transfected with GFP-FMR1 constructs from (**A**), shown as in Fig. 1A. All cells with WT-FMR1 contain the network and most cells with mutant FMR1 show network disruption (see Fig. S4C for quantification). Representative images are shown. Scale bar, 5 µm. **C.** Oligo(dT) pulldown, performed without cross-linking, of mRNA-bound FMR1 in FXR1/FXR2/FMR1 triple KO U2OS cells after ectopic expression of GFP or GFP-FMR1 constructs from (**A**). The endogenous RNA-binding protein HuR was used as positive and loading control for oligo(dT)-bound proteins. 1% of input was loaded. **D.** Live cell confocal imaging of HeLa cells transfected with GFP-FXR1-WT or FXS mutant constructs after knockdown of endogenous FXR1, shown as in (**B**). The FXS mutations G266E and I304N are located at the same amino acid positions in FXR1 and FMR1. The network is disrupted in all cells (see Fig. S4C for quantification). Representative images are shown. **E.** As in (**C**), but oligo(dT) pulldown was performed after ectopic expression of GFP-FXR1-WT, -G266E, or -I304N. **F.** Sanger sequencing results of heterozygous and homozygous N202S CC1-disrupting point mutations in endogenous *FXR1* in A549 clonal cells generated using base editing. **G.** Oligo(dT) pulldown of mRNA-bound FXR1 in A549 clonal cells from (**F**). Endogenous HuR was used as positive and loading control for oligo(dT)-bound proteins. 1% of input was loaded. **H.** Quantification of FXR1-bound mRNAs from (**G**) shown as mean ± std obtained from three independent experiments. One-way ANOVA, *** *P*<0.001. **I.** Live cell confocal imaging of A549 clonal cells from (**F**) after knockin of monomeric GFP into the endogenous *FXR1* locus. Scale bar, 5 µm. **J.** Sanger sequencing results of heterozygous KH1 domain point mutation G266E in endogenous *FXR1* in A549 clonal cells generated using prime editing. **K.** Oligo(dT) pulldown of mRNA-bound FXR1 in A549 clonal cells from (**J**). Endogenous HuR was used as positive and loading control for oligo(dT)-bound proteins. 0.2% of input was loaded. **L.** Quantification of FXR1-bound mRNAs from (**K**) shown as mean ± std obtained from three independent experiments. One-way ANOVA, *, *P*<0.05, **, *P*<0.01. **M.** Schematic summarizing the results from (**F**) to (**L**).

Epigenetic silencing of *FMR1* causes FXS^13^. In a few patients, however, single FMR1 point mutations in the KH1 (G266E) or KH2 (I304N) RNA-binding domains cause severe FXS disease symptoms^42,43^. We modeled these mutations in FMR1 and FXR1 and observed that both point mutations disrupted network assembly and reduced mRNA binding of FMR1 and FXR1 (Fig. 3B-E, S4C, S4L, Video S6 and S10)^42–44^. These results show that FMR1 also forms an mRNP network and that RNA binding is required for network assembly, suggesting that FXR1 and FMR1 need to be assembled into their respective networks to be functional.

### Single point mutations prevent assembly of the endogenous FXR1 network

To study the effects of network disruption of endogenous FXR1, we used base editing to introduce a single CC-breaking point mutation into endogenous *FXR1* in A549 cells. As only CC1 was amenable to base editing, we generated cells with an N202S mutation in FXR1 (Fig. 3F). This mutation disrupted the endogenous FXR1 network and reduced mRNA binding in oligo(dT) pulldown experiments (Fig. 3F-I).

As the CC mutation disrupts mRNA binding and FXR1 dimerization, we tested whether disruption of mRNA binding alone is sufficient to prevent FXR1 network assembly. Using prime editing, we generated the FXS patient-derived mutation G266E in the KH1 domain of endogenous *FXR1* in A549 cells (Fig. 3J). Endogenous FXR1-G266E has a strongly reduced mRNA binding ability in oligo(dT) pulldown experiments (Fig. 3K, 3L). These results reveal that disruption of mRNA binding of FXR1 is sufficient to disrupt FXR1 network assembly (Fig. 3M), indicating that mRNA is the underlying scaffold of the FXR1 network.

### Exceptionally long mRNAs bound to FXR1 dimers serve as scaffold of the FXR1 network

To start to address a potential function of the FXR1 network, we used individual-nucleotide resolution UV-cross-linking and immunoprecipitation (iCLIP) to identify FXR1-bound mRNAs in HeLa cells. To identify FXR1 network-dependent mRNAs, we depleted endogenous FXR1 and replaced it with either GFP-tagged WT or the assembly-deficient CC2-mutant FXR1 (Fig. S5A, S5B)^45^. We observed that, within mRNAs, FXR1 binds nearly exclusively to 3′UTRs or coding sequences (Fig. S5C, 50.6% and 46.8% of binding sites, respectively). We regard 2,327 mRNAs as FXR1 targets and validated 19/20 using RNA-IP followed by qRT-PCR (Fig. S5D, S5E).

We define network-dependent mRNAs as FXR1 targets whose binding is reduced by at least two-fold, when comparing the binding pattern of WT and assembly-deficient FXR1. Approximately half (*N* = 1223) of the FXR1 targets are network-dependent, whereas RNA-binding of the other half of FXR1 targets (*N* = 1104, 47%) was not affected by the assembly-deficient FXR1 mutant, and therefore are called network-independent targets (Fig. S5F, Table S1).

Comparison of network-dependent and -independent mRNAs revealed that the former have nearly twice as many FXR1 binding sites and are significantly longer, thus representing exceptionally long mRNAs with a median length of ∼6,000 nucleotides (Fig. 4A, 4B). They are also characterized by the highest AU-content and the longest 3′UTRs (Fig. 4C, S5G). Taken together, these results suggest a model whereby FXR1 dimers bind to the longest mRNAs expressed in cells, which allows them to be packaged into the FXR1 network, where they form the underlying mRNA-FXR1 dimer scaffold. Therefore, we call the network-dependent targets scaffold mRNAs of the FXR1 network. As network-independent mRNAs were only detected after cross-linking, these results suggest that they are not packaged into the network but may only associate with it. This model is consistent with the oligo(dT) pull-down experiments (see Fig. 2E), which were performed without cross-linking and only detected mRNAs strongly bound to FXR1 dimers (Fig. 2E, 3E, 3H, 3L).

**Figure 4.**
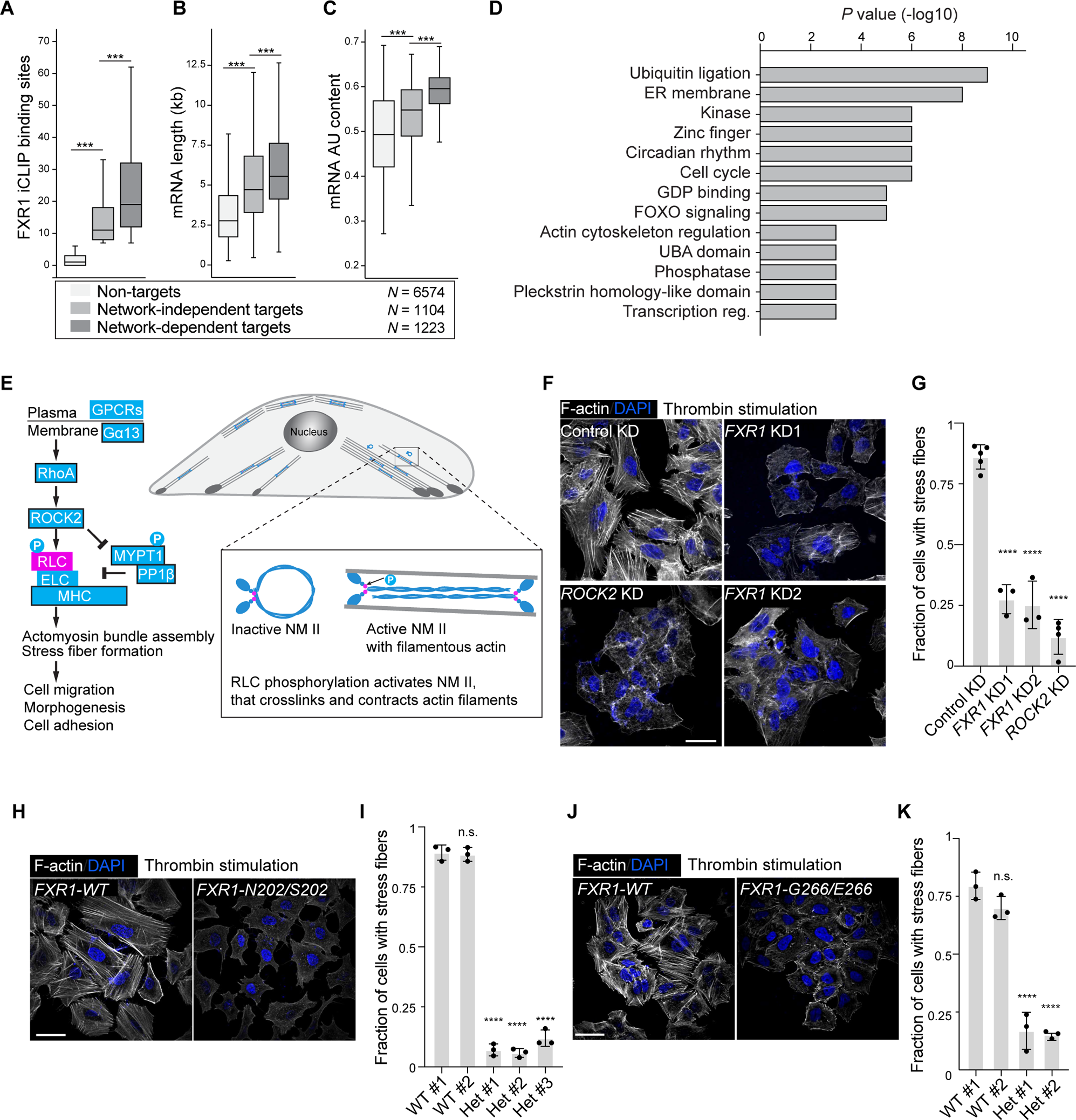
The FXR1 network is required for RhoA signaling-induced actomyosin reorganization. **A.** All mRNAs expressed in HeLa cells are grouped based on their FXR1 binding pattern. mRNAs not bound by FXR1 (*N*=6574), bound by FXR1 but network-independent (*N*=1104), bound by FXR1 and network-dependent (*N*=1223). Boxes represent median, 25^th^ and 75^th^ percentiles, error bars represent 5-95% confidence intervals. Mann-Whitney test, ***, *P<*10^−53^. **B.** As in (**A**), but mRNA length is shown. Mann-Whitney test, ***, *P<*10^−14^. **C.** As in (**A**), but AU-content of mRNAs is shown. Mann-Whitney test, ***, *P<*10^−54^. **D.** Gene ontology analysis for FXR1 network-dependent mRNA targets. Shown are the top functional gene classes and their Bonferroni-corrected *P* values. **E.** Schematic of RhoA signaling pathway-induced actomyosin remodeling. The critical signaling event for actomyosin dynamics is RLC phosphorylation of NM II. Protein symbols with black outlines are FXR1 mRNA targets. ELC, essential light chain. P, phosphorylated residue. **F.** Phalloidin staining of filamentous actin in A549 cells expressing the indicated shRNAs after serum starvation and stimulation with thrombin for 30 minutes. DAPI staining visualizes the nucleus. Representative images are shown. Scale bar, 40 µm. **G.** Quantification of the experiment in (**F**) shown as mean ± std obtained from at least three independent experiments. For each experiment and each sample at least 150 cells were counted, except for the *ROCK2* KD experiment, where 34 cells were counted. One-way ANOVA, ****, *P*<0.0001. **H.** As in (**F**), but A549 clonal cells with heterozygous N202S mutations in endogenous *FXR1* were used. Shown are representative images. **I.** Quantification of the experiment in (**H**) shown as mean ± std obtained from at least three independent experiments. For each experiment and each sample at least 28 cells were counted. One-way ANOVA, ****, *P*<0.0001. **J.** As in (**F**), but A549 clonal cells with heterozygous G266E mutations in endogenous *FXR1* were used. Shown are representative images. **K.** Quantification of the experiment in (**J**) shown as mean ± std obtained from three independent experiments. For each experiment and each sample, at least 70 cells were counted. One-way ANOVA, ****, *P*<0.0001.

### The FXR1 network provides a signaling scaffold for RhoA-induced actomyosin reorganization

To obtain insights into the physiological role of the FXR1 network, gene ontology analysis was performed to identify enriched pathways among the FXR1 scaffold mRNAs^46^. We observed a significant enrichment of various signaling pathway components, including kinases, GDP-binding proteins, and regulation of the actin cytoskeleton (Fig. 4D).

A closer look into the FXR1 targets involved in actin cytoskeleton dynamics revealed that nearly all components of the RhoA-activated actomyosin remodeling pathway are encoded by FXR1-bound mRNAs (Fig. 4E, boxes with black outline, Table S1). Dynamic regulation of the actomyosin cytoskeleton is fundamental to basically all cell types and controls cell shape, adhesion, migration, and synaptic function^47–49^. The components of the RhoA signaling pathway are ubiquitously expressed and the pathway is induced by diverse extracellular signals, such as lysophosphatidic acid (LPA) or thrombin, which activate G protein-coupled receptors, thus activating the RhoA GTPase^23,50^. Active RhoA binds and activates the Rho-associated kinase ROCK, the central regulator of actomyosin remodeling^51^. The crucial regulatory event for actomyosin remodeling is the phosphorylation of the regulatory light chains (RLC) of non-muscle myosin II (NM II). NM II is a hexamer that consists of two myosin heavy chains, two essential light chains and two RLCs. The RLCs are directly phosphorylated by ROCK^52^. RLC phosphorylation can also be increased through inhibition of phosphatase 1, which is mediated by ROCK-dependent phosphorylation of MYPT1, the regulatory subunit of phosphatase 1 (Fig. 4E). Importantly, RLC phosphorylation induces actin bundling and contraction of actin fibers, which can be read out as stress fiber formation.

To determine whether FXR1 is required for stress fiber formation, we treated A549 cells with thrombin or LPA and stained them for filamentous actin (F-actin) (Fig. 4F). RhoA stimulation-induced stress fibers were generated in cells that express control shRNAs, but their formation was strongly reduced in cells treated with shRNAs against *GNA13*, *ROCK2,* or *FXR1* (Fig. 4F, 4G, S6A-E). Since ROCK2 knockdown was sufficient to disrupt stress fiber formation and *ROCK2* mRNA was a validated FXR1 target (Fig. S5E), we focused on ROCK2 instead of ROCK1 for the rest of the study. Regulation of stress fiber formation was specific to FXR1, as FMR1 knockdown did not affect their formation (Fig. S6A-C). These results show that FXR1 protein is required for Rho A signaling-induced actomyosin remodeling. Importantly, the network-disrupting point mutations in endogenous FXR1 (N202S or G266E) also prevented stress fiber formation (Fig. 4H-K), demonstrating that not only the presence of FXR1 protein, but FXR1 assembled into the FXR1 network, is essential for RhoA signaling-induced actomyosin remodeling.

Actomyosin remodeling can positively or negatively affect cell migration^53–55^. We observed that FXR1 knockdown or ROCK inhibition impaired migration of A549 cells (Fig. S6F). When testing whether the FXR1 network is required for migration, we observed that migration in all single cell clones with WT phenotype was strongly reduced (Fig. S6G), indicating that the generation of single cell clones impairs the migration capacity of the cells, which confounded the investigation.

### Phosphorylation of RLC by ROCK2 kinase is FXR1 network-dependent

How does the FXR1 network regulate actomyosin dynamics? As FXR1 was reported to regulate translation^21^, we hypothesized that protein levels in the RhoA signaling pathway are regulated by FXR1. To identify FXR1-dependent protein abundance changes, we performed Tandem Mass Tag quantitative proteomics analysis in control and FXR1 knockdown cells. Surprisingly, among 7,067 expressed proteins, only six significantly changed expression in the absence of FXR1, and none of them were components of the RhoA signaling pathway (Fig. 5A, Table S2). Moreover, immunoblot analysis on the RhoA pathway components in unstimulated and stimulated A549 cells, in the presence or absence of FXR1, did not detect FXR1-dependent abundance changes of ROCK2, MYPT1, and the NM II subunits NM IIA and RLC (encoded by MYH9 and MYL9) (Fig. S6H-J). These results indicated that FXR1 does not widely affect protein abundance in the investigated cell types and does not control protein levels of the RhoA signaling pathway.

**Figure 5.**
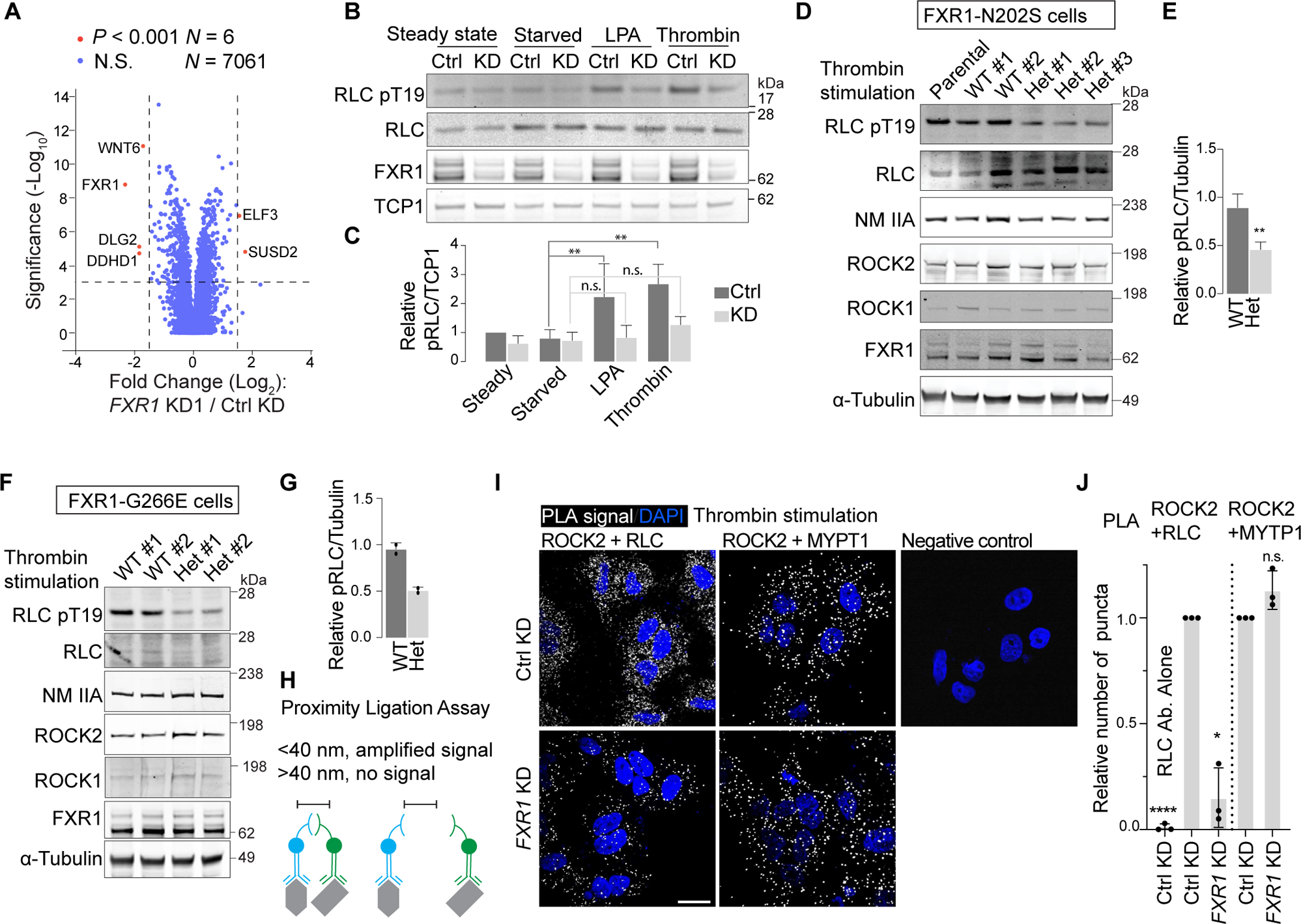
Phosphorylation of RLC by ROCK2 kinase is FXR1 network dependent. **A.** Tandem Mass Tag quantitative proteomics analysis of HeLa cells after control or *FXR1* KD. Proteins whose abundance was significantly affected by *FXR1* KD are colored red (*N*=6), whereas proteins not significantly affected are colored in blue (*N*=7061). **B.** Western blot of the indicated endogenous proteins in A549 cells grown in the indicated conditions. Ctrl, expressing control shRNA, KD, expressing FXR1 shRNA1. TCP1 was used as loading control. **C.** Quantification of phospho-RLC level from (**B**) shown as mean ± std obtained from at least three independent experiments. One-way ANOVA, **, *P*< 0.01. n.s., not significant. **D.** Western blot of the indicated proteins in serum-starved and thrombin-stimulated parental A549 and clonal cell lines containing WT FXR1 or a heterozygous N202S mutation in endogenous *FXR1*. α-Tubulin was used as loading control. **E.** Quantification of phospho-RLC level from (**D**) shown as mean ± std obtained from three clonal cell lines each. One-way ANOVA, **, *P*<0.01. **F.** Western blot of the indicated proteins in serum-starved and thrombin-stimulated parental A549 and clonal cell lines containing WT FXR1 or a heterozygous G266E mutation in endogenous *FXR1*. α-Tubulin was used as loading control. **G.** Quantification of phospho-RLC level from (**F**) shown as mean ± std obtained from two clonal cell lines each. **H.** Schematic of the proximity ligation assay (PLA), which generates a positive signal if the distance between two endogenous proteins is smaller than 40 nm. **I.** PLA performed in serum-starved thrombin-stimulated A549 cells, indicating that FXR1 is required for proximity between ROCK2 and RLC, but not for proximity between ROCK2 and MYPT1. As negative control, the RLC antibody alone was used. DAPI staining visualizes the nucleus. Representative images are shown. Scale bar, 20 µm. **J.** Quantification of the experiment in (**I**), shown as mean ± std of three independent experiments. For each experiment and each sample at least 39 cells were counted. One-way ANOVA, ****, *P*<0.0001.

To identify the molecular mechanism by which the FXR1 network impacts the signaling pathway that controls actomyosin remodeling, we examined the pathway in greater detail. As FXR1 knockdown did not reduce the amount of active RhoA obtained through GPCR stimulation (Fig. S6K), we concluded that the RhoA pathway upstream of ROCK is unaffected by FXR1 deficiency. We then discovered that RhoA signaling-induced RLC phosphorylation was FXR1 dependent (Fig. 5B, 5C). Importantly, RLC phosphorylation was impaired not only in cells with knockdown of FXR1, but also impaired in cells with network-disrupting mutations (N202S or G266E) of endogenous FXR1 (Fig. 5D-G). These data indicate that the FXR1 network is essential for RhoA-signaling induced phosphorylation of NM II.

### The FXR1 network provides proximity between the ROCK2 kinase and its substrate RLC

Phosphorylation of RLC requires an active ROCK2 kinase and spatial proximity between kinase and substrate^51,56^. As phosphorylation of the ROCK2 substrate MYPT1 was FXR1-independent, we concluded that ROCK2 activation does not rely on FXR1 (Fig. S6I, S6J). To determine whether FXR1 acts as a scaffold for ROCK2 kinase and its substrate RLC, we performed a Proximity Ligation Assay (PLA) in cells expressing control or FXR1-targeting shRNAs. PLA allows the in-situ detection of protein:protein interactions whose distance is less than 40 nm (Fig. 5H)^57^. After thrombin stimulation, the ROCK2 kinase is in proximity with both its substrates MYPT1 and RLC in control cells, whereas in FXR1 knockdown cells, the proximity between ROCK2 and RLC is strongly reduced (Fig. 5I, 5J, S6L).

Taken together, these results show that FXR1 is essential for RhoA signaling-induced actomyosin remodeling, where the crucial signaling step is an FXR1 network-dependent event that establishes spatial proximity between kinase and substrate. As FXR1 has a large number of protein:protein interaction domains (Fig. 2A), we hypothesized that the FXR1 network may therefore act as signaling hub.

### Network-dependent protein interactors have similar protein domains as FXR1

To identify network-dependent protein:protein interactors of FXR1, we performed GFP co-immunoprecipitation (co-IP) and SILAC proteomics analysis using GFP-FXR1 WT and assembly-deficient CC2 mutant, expressed in cells depleted of endogenous FXR1 (Fig. 6A, S7A). We identified several proteins, including FXR2, FMR1, UBAP2L, TOP3B, TDRD3, PRRC2C, PRRC2A, and AP2A1, that interacted significantly better with WT FXR1 compared with CC2 mutant FXR1 (Fig. 6A, Table S3). To validate these results, we performed co-IP in the presence or absence of RNase A, followed by immunoblot analysis. This approach validated 10/10 candidates (Fig. 6B, 6C). We observed that most of these protein:protein interactions are RNA-dependent, which supports their FXR1 network dependence (Fig. 6B, 6C).

**Figure 6.**
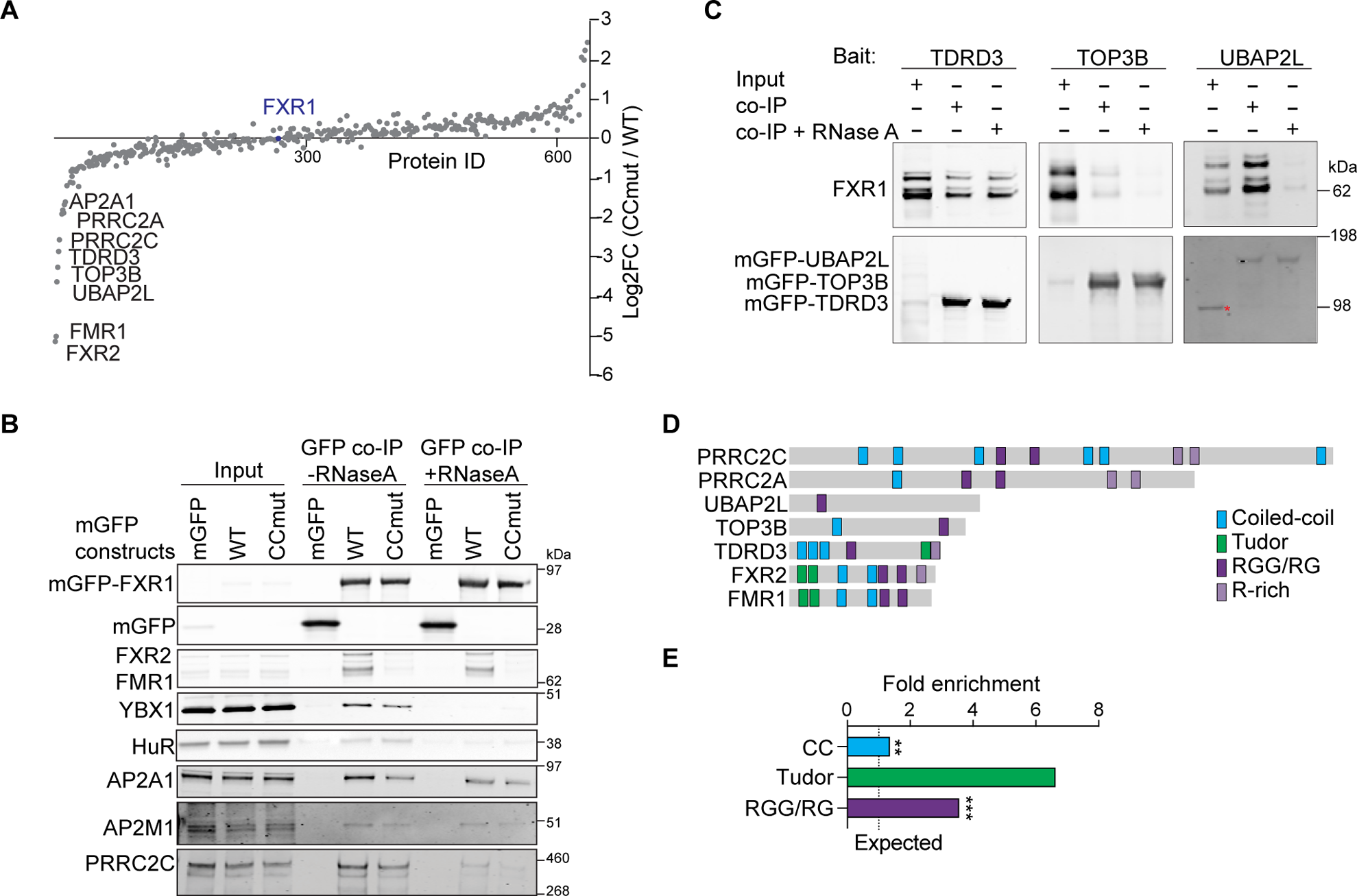
FXR1 network-dependent protein interactors contain CC, Tudor, and RGG domains. **A.** SILAC mass spectrometry analysis of HeLa cells. Shown is log2 fold change (FC) of protein counts of CC2mut/WT samples. Reduced interaction in CC2 mutant samples indicates that the interaction with FXR1 is network dependent. The top network dependent FXR1 interactors are indicated. For full list, see Table S3. **B.** Validation of the SILAC proteomics results using GFP co-IP of the indicated endogenous proteins followed by western blot in the presence or absence of RNase A. GFP-FXR1 constructs were ectopically expressed in HeLa cells depleted of endogenous FXR1. 0.5% input was loaded. **C.** As in (**B**), but GFP co-IP of endogenous FXR1 by ectopically expressed interactors. The red star symbol marks an unspecific band. 1% input was loaded. **D.** Protein domains of the top FXR1 network-dependent interactors. Shown are CC, Tudor, RG/RGG, and R-rich domains in color. **E.** Fold enrichment of indicated protein domains in the 20% of proteins from (**A**) with the most negative FC. Shown is the observed-over-expected frequency. Chi-square test, **, *P*=0.002, ***, *P*<0.0001. Chi-square test for Tudor domains cannot be performed as the numbers are too small. See Table S3.

When analyzing the protein domains of the network dependent FXR1 interactors, we made the surprising observation that the interactors contain the same kinds of protein domains as FXR1 (Fig. 6D). FXR1 contains CC, Tudor, and RGG domains and these domains were significantly enriched among the top 20% of network-dependent FXR1 binding partners (Fig. 6E, Tables S1 and S3). Moreover, FXR1 mRNA targets were significantly enriched among the FXR1 protein interactors (Table S3). As CC, Tudor and RGG domains can perform homo- and heterodimerization^28,30,37–39^, these data suggest that proteins may use these domains to become recruited into the FXR1 network, thus acting as protein clients of the network. We hypothesized that signaling proteins containing these domains become recruited into the FXR1 network and use the network to achieve spatial proximity.

### The CC domain of ROCK2 binds to FXR1

FXR1 network-dependent proximity occurs between ROCK2 and NM II (Fig. 5H-J). Both ROCK2 and NM II contain large CC domains (Fig. 7A). To determine whether the CC domains of ROCK2 interact with FXR1, we performed co-IP of GFP-tagged ROCK2 truncation constructs (Fig. 7B). We observed that the C-terminal half of ROCK2 strongly interacts with FXR1 (Fig. S7B, S7C). As the interaction requires the presence of the CC2 domain of ROCK2, the results indicate that this CC domain is necessary for FXR1 binding (Fig. 7C). This finding is consistent with a model whereby proteins that contain binding sites for FXR1 are recruited into the FXR1 network (Fig. S7D).

**Figure 7.**
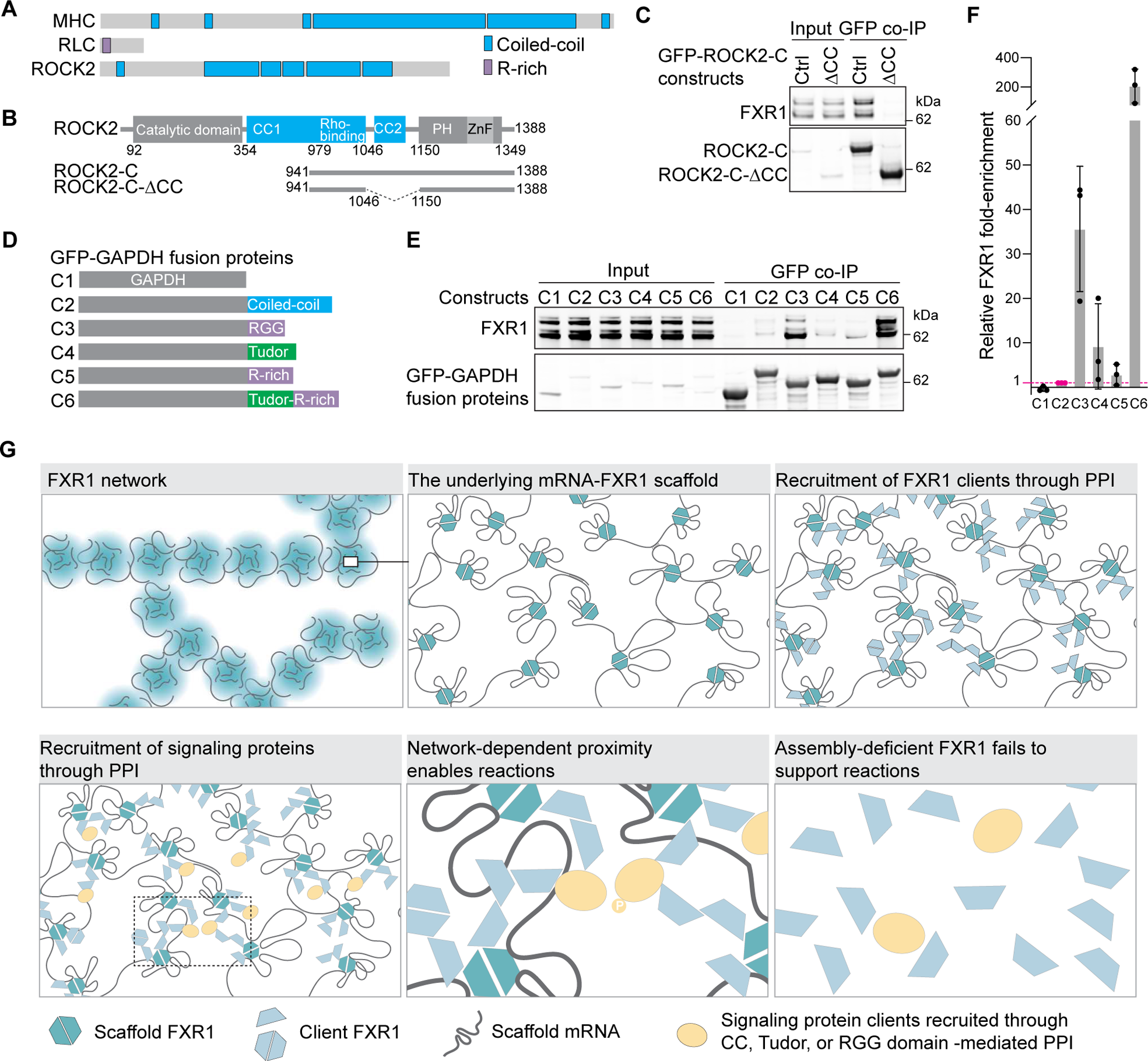
The presence of CC, Tudor, or RGG domains is sufficient for binding to FXR1. **A.** Protein domains of NM II (MYH9), RLC (MYL9), and ROCK2. Highlighted are CC and R-rich domains. **B.** Amino acid boundaries of ROCK2 protein domains and schematics of ROCK2 constructs. The numbers indicate amino acids. **C.** GFP co-IP, followed by western blot of endogenous FXR1 after ectopic expression of GFP-ROCK2-C or GFP-ROCK2-C-ΔCC (from **B**) in HeLa cells. 1% input was loaded. **D.** Schematic of GFP-GAPDH fusion constructs. The following domains were fused to GAPDH: CC2 domain of ROCK2, RGG domain of TOP3B, Tudor domain of TDRD3, R-rich region of TDRD3, and both the Tudor and R-rich regions of TDRD3. See Fig. S7E for their amino acid sequences. **E.** GFP co-IP, followed by western blot of endogenous FXR1 after ectopic expression of GFP-GAPDH fusion constructs from (**D**) in HeLa cells. A representative experiment is shown. **F.** Quantification of (**E**). Shown is FXR1 enrichment normalized to sample C2 (shown in magenta) as mean ± std obtained from at least three independent experiments. **G.** Model of the FXR1 network and its function as a scaffold for signaling reactions by establishing spatial proximity between kinases and their substrates. P, phosphorylated residue. See text for details.

### CC, Tudor or RGG domains are sufficient for binding to FXR1

Finally, we determined whether the presence of a single proposed domain (CC, Tudor, or RGG) was sufficient for binding to FXR1. GFP-tagged GAPDH, an enzyme that does not interact with FXR1, was fused to either the CC2 domain of ROCK2, the RGG domain of TOP3B, the Tudor domain of TDRD3, the R-rich domain of TDRD3, or both domains (Fig. 7D, S7E). Co-IP demonstrated that all GAPDH-fusion proteins interacted with endogenous FXR1, whereas GAPDH alone did not (Fig. 7E, 7F). Whereas the presence of a single FXR1 protein interaction domain was sufficient for FXR1 binding, the binding was weak for two of the four tested domains. Importantly, the presence of two interaction domains, such as a Tudor and an R-rich domain, showed a cooperative effect for FXR1 binding and increased the affinity by ∼20-fold (Fig. 7E, 7F).

## Discussion

Here, we report the discovery of the FXR1 network—a large cytoplasmic mRNP network that acts as a multivalent signaling platform. The FXR1 network is present throughout the cytoplasm of all cells so far investigated. In addition to spherical condensates like P bodies and stress granules, our work shows that the cytoplasm is further compartmentalized by several network-like condensates, including TIS granules and the FXR1 network^5–7^.

### Regulation of proximity of signaling proteins by the FXR1 network

The underlying scaffold of the FXR1 network are exceptionally long mRNAs that are bound and packaged by FXR1 dimers (Fig. 7G). Only a minority of FXR1 stably binds to mRNA and is part of the underlying scaffold. As FXR1 is nearly entirely present within high-molecular-weight complexes, most FXR1 molecules act as clients and are recruited into the network using protein:protein interactions through multiple CC, Tudor, and RGG domains, which are known for their homo- and heterodimerization capacities^28,30,37–39^. Homodimerization recruits FXR1 molecules into the network, whereas heterodimerization recruits other proteins, such as signaling factors. The high concentration of FXR1 molecules in the network generates a high concentration of binding sites for CC, Tudor, and RGG domains and allows multivalent binding of recruited clients, including signaling proteins, which brings these molecules into proximity with each other (Fig. 7G).

Point mutations in the KH domains or in the CC domains prevent RNA binding of FXR1 and prevent formation of the network scaffold, which results in diffusive FXR1 protein. Network disruption lowers the local FXR1 concentration, thus preventing transient trapping of signaling molecules and network-dependent spatial proximity, which impairs enzyme-substrate interactions and prevents productive signal transduction (Fig. 7G). Thus, the FXR1 network brings proteins containing certain CC, Tudor, or RGG domains into proximity to promote key signaling pathways, as we demonstrated for actomyosin remodeling. As many other signaling proteins also contain these domains^29^, it is likely that additional signaling pathways use the FXR1 network as scaffold.

### The FXR1 network is essential for actomyosin remodeling and is disrupted by disease mutations

Single point mutations (G266E or I304N) in FXR1 disrupt the FXR1 network. The mutations were detected in the FXR1 homolog FMR1, where they cause FXS^13,27,42^. FXS is the most common inherited cause of intellectual disability and is one of the most common inherited causes of ASD^20^. Variants in the *FXR1* gene are also strongly associated with increased risk for ASD and schizophrenia^14–17^. Deletion of FXR1 in mouse interneurons reduces their excitability and causes schizophrenia-like symptoms^20^, suggesting a role for FXR1 in neuronal functions.

One of the physiological phenotypes caused by FXR1 network disruption is impaired actomyosin cytoskeleton remodeling, a process that occurs in nearly all cell types^47–49^. In non-neuronal cells, it is essential for the regulation of cell shape, adhesion, migration, and tissue architecture, whereas in neuronal cells it also controls dendritic spine morphology and synaptic function^47–49^. Alterations in spine morphology are associated with neuronal dysfunction and can lead to cognitive and behavioral problems^58,59^. Therefore, we suggest that FXR1 network disruption, which impairs actomyosin dynamics, could be one of the underlying causes of abnormal dendritic spine morphology and synaptic function in patients with FXS.

### Do FXR1 and FMR1 have overlapping functions?

FMR1 is also present as mRNP network in the cytoplasm. Moreover, FXR family members bind to each other and are incorporated into each other’s networks^26^. To address whether FXR1 and FMR1 have overlapping functions, we tested the requirement of FMR1 for stress fiber formation and found that in A549 cells, only FXR1 was necessary for RhoA signaling-induced stress fiber formation. We suspect that the functions of FXR family proteins strongly depend on their expression levels, because dosage reduction of assembly-competent WT FXR1 in the samples with heterozygous FXR1 mutations was sufficient to impair stress fiber formation. The mRNA expression pattern of the three FXR family homologs shows that FXR1 is expressed ubiquitously at very high levels, whereas FMR1 and FXR2 are mostly expressed in the brain, suggesting that in non-neuronal cell types FXR1’s function may be dominant.

### Molecular principles of the mRNA scaffold

FXR1 binds and packages the longest ∼1200 mRNAs expressed in cells, which results in the formation of an mRNP network. Most mRNAs are packaged into individual mRNPs by the exon-junction complex, which binds to exon-intron junctions in coding sequences^60^. The FXR1-bound mRNAs have very long 3′UTRs, which lack exon-intron junctions, suggesting that FXR1 may have a packaging function for these mRNAs. This idea is supported by the ubiquitous and high expression of FXR1^23^, which suggests that the role of FXR1 is required in all cells. Moreover, the intrinsic binding affinity of FXR1 to RNA seems very weak^61^. We speculate that the weak RNA binding affinity of FXR1 is responsible for the selection of long 3′UTRs as they provide the largest number of potential binding sites.

In addition to the FXR1 network, also TIS granules have a network-like structure, generated through RNA-RNA interactions^10^. We showed that FXR1 network formation requires RNA binding of the RG motif in the FXR1 IDR. RG motifs bind to RNA and remodel RNA-RNA interactions during RNA annealing reactions^36,62,63^, suggesting that RG motifs play crucial roles in the formation of network-like condensates.

### Molecular principles of FXR1 network-dependent proximity of clients

We showed how a large mRNP network serves as signaling scaffold for proteins that do not contain RNA-binding domains. Specificity of the FXR1 network-based signaling platform is provided by CC, Tudor, and RGG domains. In addition to Tudor-Tudor or RGG-RGG interactions, Tudor-RGG interactions are also possible, as Tudor domains bind to methylated arginines, usually in the context of RG/RGG domains^25,29^. RG/RGG domains seem to be the most versatile domains in this system as they can bind to RNA and protein^25,28–30^. Although RGG domains are often found in nuclear and RNA-binding proteins, in the cytoplasm, they are observed in structural and regulatory factors, including intermediate filaments, cytoskeleton-binding proteins, and kinases^29^. Therefore, we propose that cytoskeletal processes that need to be coordinated within the entire cytoplasm may take advantage of the FXR1 network because it provides a scaffold to promote signaling events throughout the cytoplasm.

In addition to proteins or lipid membranes acting as signaling scaffolds^64,65^, we uncovered another type of signaling scaffold in the form of an mRNP network. Its underlying scaffold is generated by FXR1-bound mRNAs, revealing that mRNAs perform structural roles in the cytoplasm. We show that the function of mRNAs and RNA-binding proteins can go beyond the regulation of mRNA-based processes^66^. So far, RNA-binding proteins are generally considered to regulate mRNA stability, translation, or localization. However, we demonstrate that they can affect signaling pathways and cytoskeleton processes, thus broadening the impact of mRNA and RNA-binding proteins on cellular processes.

### Limitations of the study

This study was performed with cell lines grown in culture. Therefore, the physiological functions of the FXR1 network in living animals are currently unknown. We documented the requirement of the FXR1 network for one step of an important signaling pathway. However, the FXR1 mRNA targets are enriched for many other signaling factors, including ubiquitin ligases, but we currently do not know the scope of signaling reactions that are FXR1 network dependent. To identify FXR1 network-dependent interactors, we used affinity purification-mass spectrometry, but this method only captures interacting proteins with relatively high affinity or abundance. During cell lysis, protein concentration is strongly reduced, potentially leading to the loss of low affinity interactors. This is relevant for the study of condensates, where protein concentration is key to condensate formation. Labeling the neighboring molecules before cell lysis through proximity ligation may provide more FXR1 network-enriched signaling proteins to allow identification of additional FXR1 network-dependent biochemical reactions.

## Acknowledgements

This work was funded by the NIH Director′s Pioneer Award (DP1-GM123454), the NIH grant R35GM144046, a grant from the Pershing Square Sohn Cancer Research Alliance, and the MSK Core Grant (P30 CA008748) to CM. J.U. received funding from the ERC under the European Union Horizon 2020 Research and Innovation Program (835300-RNPdynamics). X.C. received the Kravis Women in Science Endeavor (WiSE) post-doctoral fellowship. We thank the Molecular Cytology Core Facility and its staff for support in confocal microscopy. The facility is funded by MSK Core Grant (P30 CA008748).

## Author contribution

X.C. performed all experiments, except the FXR1 iCLIP experiment, which was performed and analyzed by U.J. and J.U. M.M.F provided the gene features. X.C. and C.M. conceived the project, designed the experiments, and wrote the manuscript with input from all authors. We thank members of the Mayr lab, and Nancy Bonini (U Penn) for critical reading of the manuscript and for helpful discussions.

## Supplementary Figure Legends

**Figure S1.**
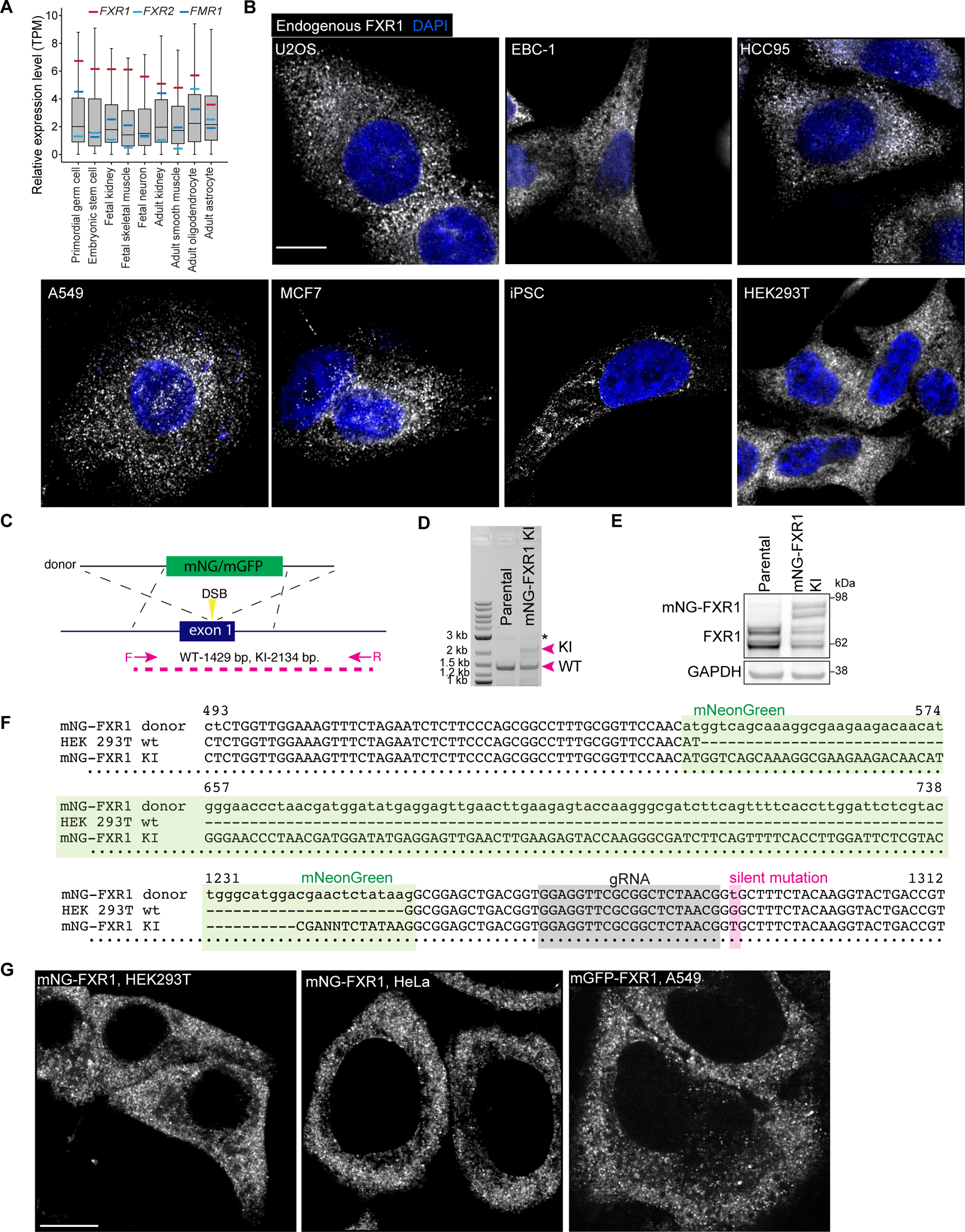
Gene and protein expression pattern of endogenous FXR1, related to Figure 1. **A.** The gene expression level of the FXR family proteins in various primary cells and tissues. The red, blue, and light blue bars represent the mRNA expression levels of FXR1, FMR1, and FXR2, respectively. The boxplots show the distribution of expression levels of all expressed mRNAs in the indicated cell types, obtained from Han et al., (2020)^23^. Boxplots shown as in Fig. 4A. **B.** Representative confocal images of immunofluorescence staining of endogenous FXR1 proteins in the indicated cell lines. U2OS, human osteosarcoma epithelial cell line; EBC-1, HCC95, A549 are human lung carcinoma lines; MCF7, human breast cancer line; iPSC, human induced pluripotent stem cells; HEK293T, human immortalized embryonic kidney cells. **C.** Knockin strategy of mGFP or mNG into the N-terminus of endogenous *FXR1* using a CRISPR-based approach. **D.** Genotyping agarose gel with primer pairs shown in (**C**). The black star symbol marks an unspecific PCR product. **E.** Western blotting of FXR1 in parental and mNG knockin HEK293T cell lines. **F.** Sanger sequencing results of the two PCR bands marked with magenta arrows in (**D**), aligned to the mNG-FXR1 donor sequence. mNG, gRNA, and the introduced silent mutations are highlighted with green, gray, and magenta boxes, respectively. **G.** Live cell confocal imaging of endogenous FXR1 tagged with either mNG or mGFP in the indicated cell lines. Representative images are shown. Scale bar, 10 µm.

**Figure S2.**
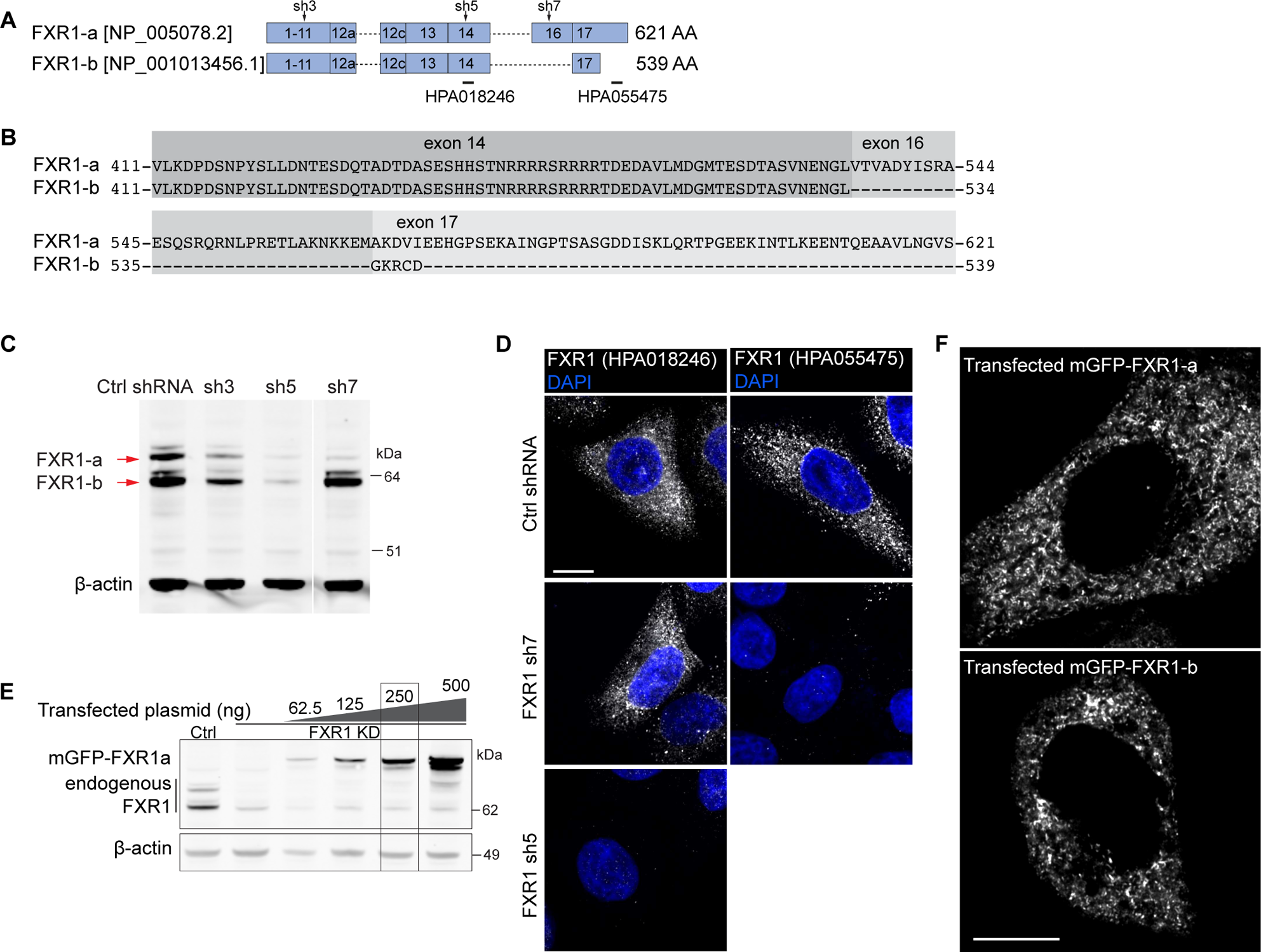
All main FXR1 isoforms can form the FXR1 network, related to Figure 1. **A.** Gene model depicting the exon structure of the two most common FXR1 splice isoforms in non-muscle cells. The three shRNAs targeting FXR1 exons used in this study are highlighted as sh3, sh5, and sh7. The epitope locations of the two antibodies used for immunofluorescence staining shown in **(D)** are labeled. **B.** The sequences of the C-terminal ends of human FXR1 isoforms a and b are shown. **C.** Western blot of the indicated endogenous FXR1 proteins in HeLa cells stably expressing a control (ctrl) shRNA (targeting luciferase) or shRNA3, shRNA5, and shRNA7 against FXR1. **D.** Immunofluorescence staining of endogenous FXR1 protein in HeLa cells expressing the control shRNA or the indicated FXR1-targeting shRNAs from (**A**). Isoform-specific antibodies, as indicated in (**A**) were used. All cells contain the network and representative confocal images are shown. Scale bar, 10 µm. **E.** Western blot of FXR1 in HeLa cells expressing control shRNA or FXR1-targeting shRNA5 transfected with increasing amounts of shRNA5-resistant mGFP-FXR1 constructs. The boxed condition was used for the rest of the study. **F.** Live cell confocal imaging of HeLa cells transfected with the indicated FXR1 constructs. Representative images are shown. See Fig. S4C for quantifications. Scale bar, 10 µm.

**Figure S3.**
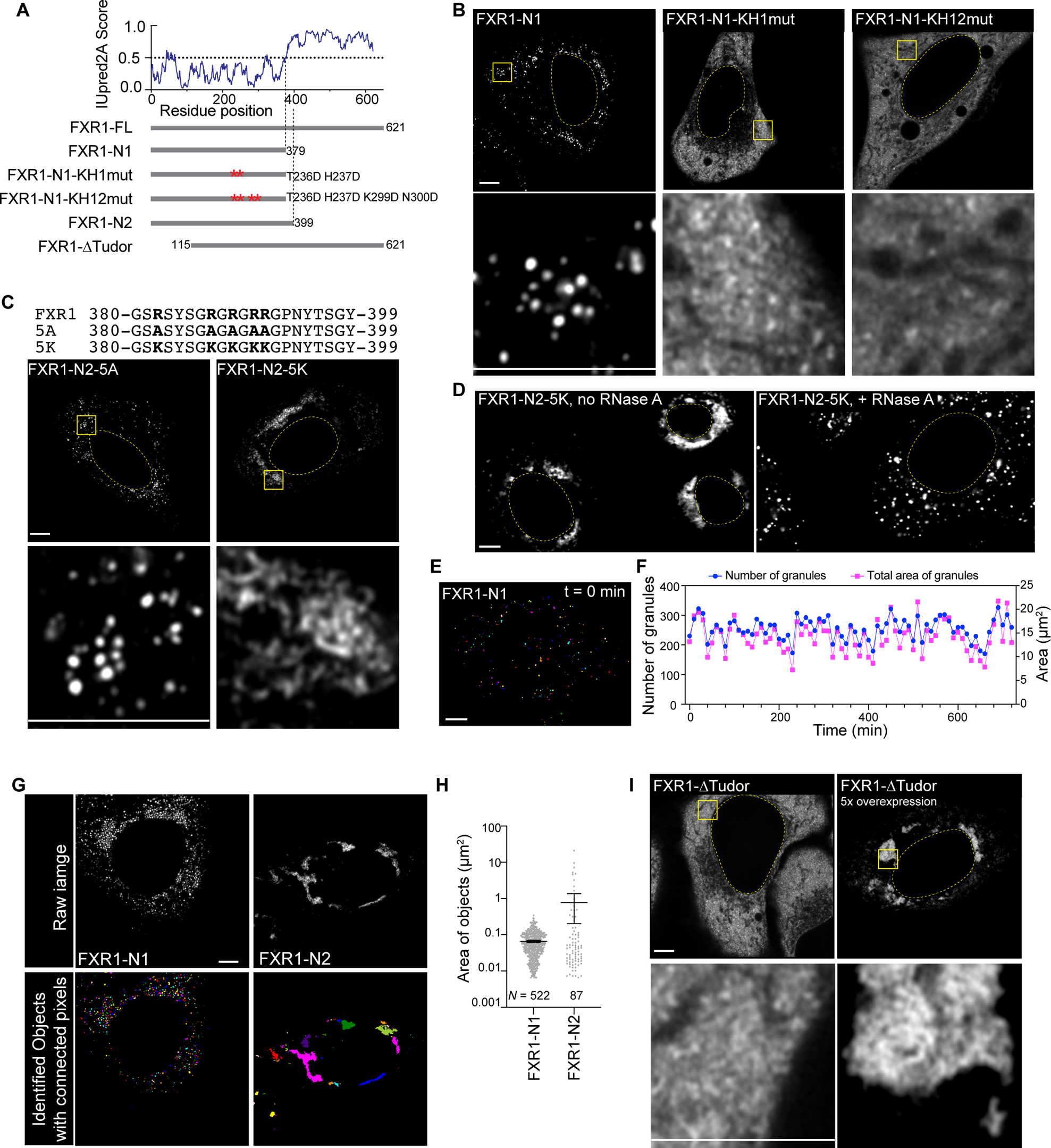
Formation of FXR1 granules and the FXR1 network requires RNA, related to Figure 1. **A.** Human FXR1 IUpred2A score and schematics of the used constructs. The GXXG motif, required for RNA binding of FXR1 KH domains was mutated to GDDG. Red star symbols represent the positions of the introduced mutations. **B.** Live cell confocal imaging of HeLa cells depleted of endogenous FXR1 and transfected with the indicated constructs. Representative images are shown as in Fig. 1A. See Fig. S4C for quantifications. Scale bar, 5 µm. **C.** Live cell confocal imaging of HeLa cells depleted of endogenous FXR1 and transfected with the indicated constructs, shown as in Fig. 1A. All cells expressing FXR1-N2-5A generated spherical granules, whereas all cells expressing FXR1-N2-5K generated a network. Representative images are shown. See Fig. S4C for quantification. The 20 aa sequence that distinguishes FXR1-N2 from FXR1-N1 is shown and the arginine residues that are mutated are shown in bold. Scale bar, 5 µm. **D.** Confocal imaging of HeLa cells transfected with GFP-FXR1-N2-5K after digitonin permeabilization in the presence or absence of RNase A treatment for 30 minutes. Representative images from at least three independent experiments are shown, where 40 cells were examined. Scale bar, 5 µm. **E.** Frame one from the timelapse of GFP-FXR1-N1 analyzed in (**F**). The timelapse was recorded at an interval of 10 min, spanning 12 hours. Scale bar, 5 µm. **F.** Quantification of the number and total area of the granules from the timelapse shown in (**E**). The fluctuations represent the granules entering and leaving the imaging plane. **G.** Confocal imaging of HeLa cells transfected with GFP-FXR1-N1 or -N2 and their corresponding identified objects with connected pixels randomly colored. Scale bar, 5 µm. **H.** Quantification of the object size shown as area (µm^2^) from the images in (**G**). The data is presented as mean ± 95% CI. The number of objects identified is 522 and 87 for FXR1-N1 and -N2, respectively.

**Figure S4.**
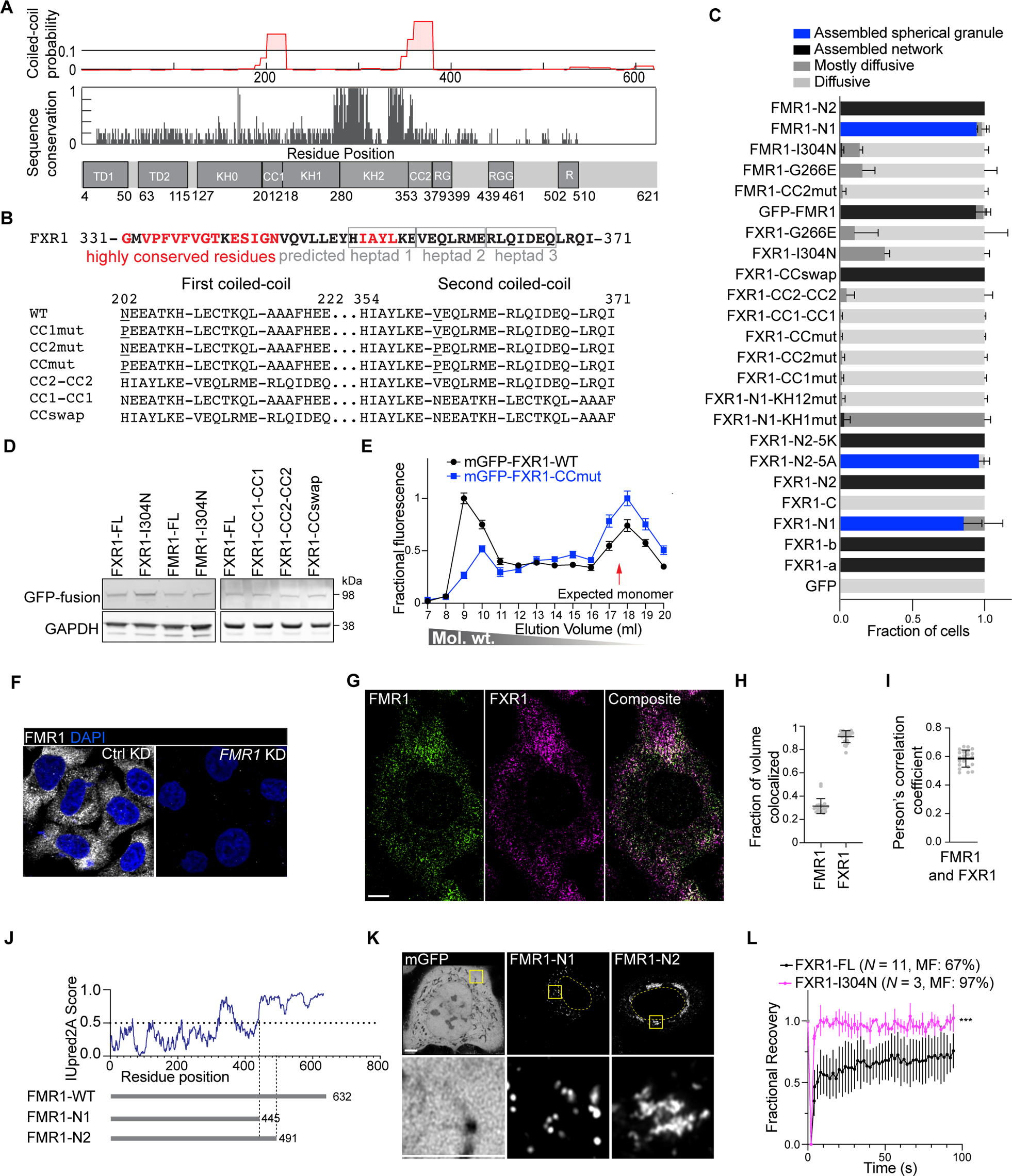
FMR1 assembles into an mRNP network using the same principles as identified for FXR1 and details on FXR1 CC mutants are shown, related to Figures 2 and 3. **A.** Human FXR1 protein domain boundaries and amino acid (aa) sequence conservation score across metazoa. Also shown is the probability for CC formation according to NCOILs. **B.** The three heptads in the predicted FXR1 CC2 domain and their neighboring aa are shown. Highly conserved residues from (**A**) are shown in red. The aa sequences of the FXR1 CC mutant constructs are shown in the bottom panel. The first heptad of CC2 was not targeted in any of the mutants because of its high conservation score. **C.** Quantification of GFP-FXR1 or GFP-FMR1 signal distribution pattern of transfected fusion constructs used in this study. A total of at least 53 cells from three or more independent experiments were scored and shown as mean ± std. The GFP signal was scored as diffusive, mostly diffusive (as shown in Fig. S3B, FXR1-N1-KH1mut), assembled network, or spherical granule. **D.** Western blot of ectopically expressed GFP-fusion proteins show comparable expression levels across samples. GAPDH was used as loading control. **E.** Size exclusion chromatography of cells shown in Fig. 2C. GFP-FXR1 fluorescence was measured using a plate reader. Shown is mean ± std of three technical replicates obtained from one fractionation experiment. **F.** Immunofluorescence staining of endogenous FMR1 protein in HeLa cells expressing the control shRNA and FMR1-targeting shRNAs. The antibody used for immunofluorescence staining was clone 6B8 (BioLegend, Cat# 834601). Scale bar, 20 µm. **G.** Representative deconvolved images of FMR1 (green) and FXR1 (magenta) double immunofluorescence staining in HeLa cells. Scale bar, 5 µm. **H.** Quantification of the fraction of colocalized volumes for FXR1 and FMR1 shown as mean ± std from 21 high-resolution volumes of HeLa cells. **I.** Pearson’s correlation coefficient between FXR1 and FMR1 fluorescence signals shown as mean ± std quantified from 21 high-resolution HeLa cells. **J.** Human FMR1 IUpred2A score and schematics of the used FMR1 constructs. **K.** Live cell confocal imaging of HeLa cells expressing the indicated GFP-FMR1 constructs. Representative images are shown as in Fig. 1A. See Fig. S4C for quantifications. Scale bar, 5 µm. **L.** FRAP analysis of GFP-FXR1-FL and -I304N expressed in HeLa cells. Shown is the normalized FRAP curve as mean ± std from at least three cells each. MF: mobile fraction. See Videos S7 and S10 for representative fluorescence recovery. Mann-Whitney test, ***, *P<*10^−57^.

**Figure S5.**
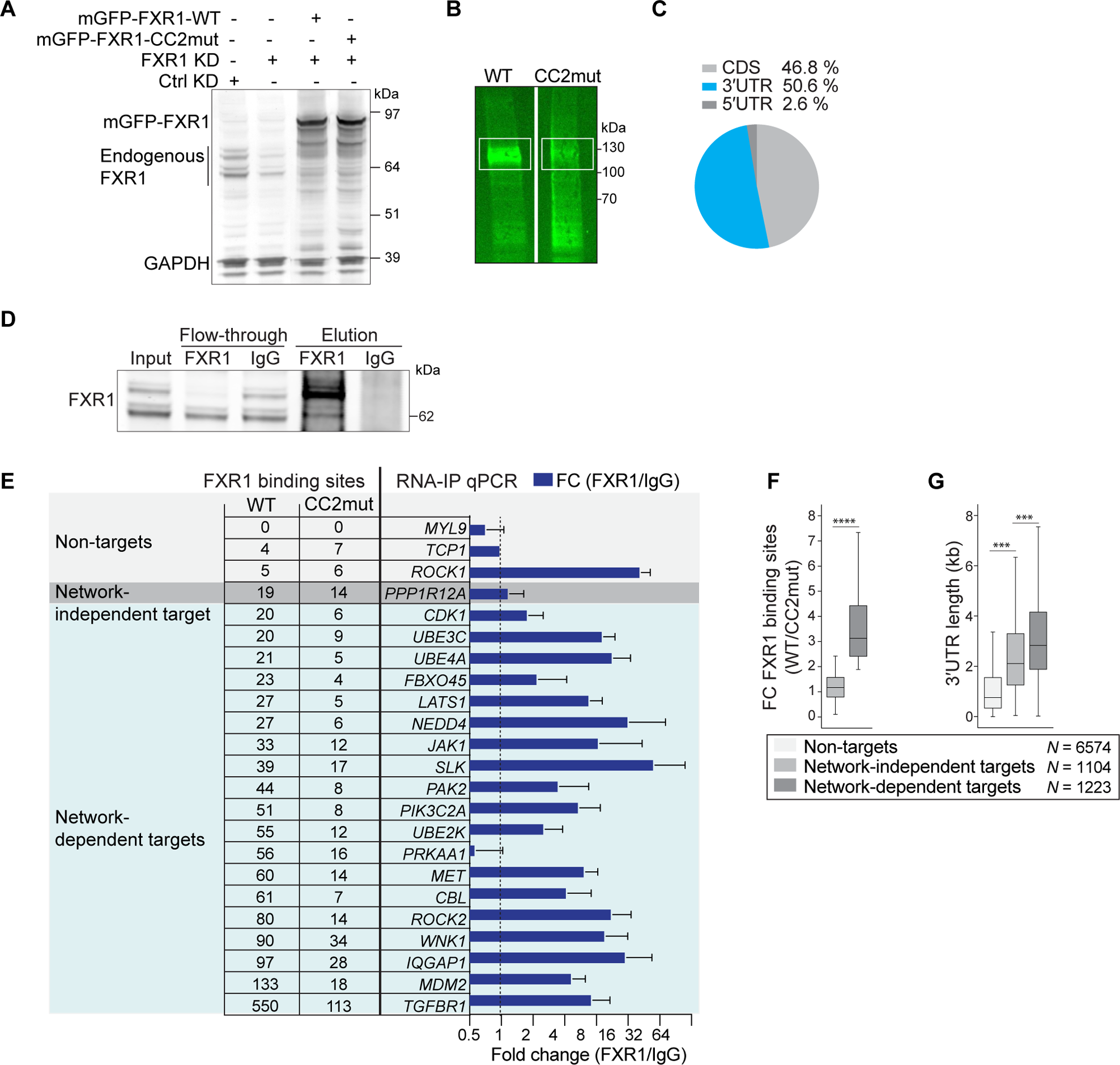
Identification of FXR1 network assembly-dependent mRNA targets using iCLIP and their validation, related to Figure 4. **A.** Western blot of endogenous and transfected FXR1 in HeLa cells expressing control shRNA or FXR1-targeting shRNA5, transfected with shRNA5-resistant mGFP-FXR1-WT or mGFP-FXR1-CC2mut. The samples in lanes 3 and 4 were crosslinked for the iCLIP experiment. GAPDH was blotted as loading control. **B.** Infrared scan showing crosslinked RNA and FXR1 complexes separated by SDS-PAGE. The boxed regions were isolated for iCLIP sample preparation. **C.** Pie chart showing the genomic distribution of unique iCLIP reads for FXR1 in CDS, 5′UTR, and 3′UTRs. **D.** Western blot of endogenous FXR1 with samples used in RNA immunoprecipitation (RIP) without cross-linking. The FXR1 antibody (Novus Biologicals, NBP2-22246) predominantly enriched FXR1 isoform a, whereas IgG did not enrich any FXR1 protein. **E.** The number of FXR1 binding sites found in specified mRNAs is shown on the left. The right part of the panel shows the fold change in RNA-immunoprecipitation (RIP) signal obtained without cross-linking using FXR1 antibody compared to IgG, obtained by RT-qPCR analysis of the indicated mRNAs in HeLa cells. Shown is mean ± std of three independent experiments. **F.** Identification of network-dependent (*N* = 1223) and network-independent (*N* = 1104) FXR1 mRNA targets. Network-dependent targets were defined based on a reduction of at least two-fold in FXR1 binding sites observed by iCLIP, when comparing WT and CC2mut FXR1. Boxplots are shown as in Fig. 4A. Mann-Whitney test, ****, *P* = 0. **G.** Distribution of 3′UTR length in the three groups from Fig. 4A and shown as in Fig. 4A. Mann-Whitney test, ***, *P <* 10^−25^.

**Figure S6.**
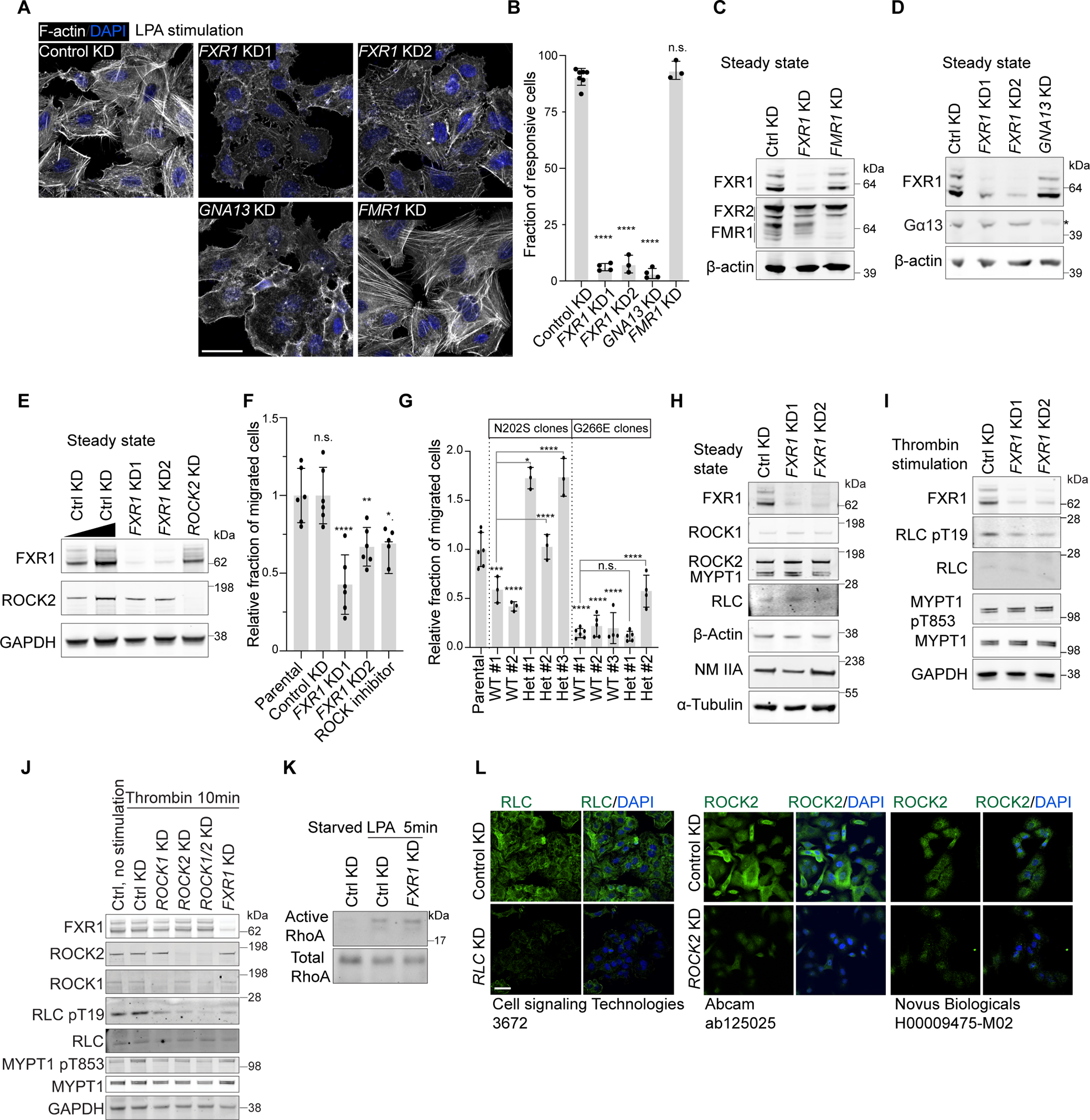
FXR1-dependent regulation of the RhoA signaling pathway, related to Figure 4 and Figure 5. **A.** Phalloidin staining of filamentous actin in A549 cells expressing the indicated shRNAs after serum starvation and stimulation with LPA for 30 minutes. DAPI staining visualizes the nucleus. Representative images are shown. Scale bar, 40 µm. **B.** Quantification of the experiment in (**A**) shown as mean ± std obtained from at least three independent experiments. For each experiment and each sample at least 92 cells were counted. One-way ANOVA, **** *P* < 0.0001. n.s., not significant. **C.** Western blot of the indicated endogenous proteins from A549 cells shows knockdown efficiency of shRNAs targeting *FXR1* and *FMR1*. The knockdown was specific, as no cross-effect on FXR family proteins was observed. **D.** As in (**C**), but knockdown efficiency of shRNAs targeting *FXR1* and *GNA13* is shown. **E.** As in (**C**), but knockdown efficiency of shRNAs targeting *FXR1* and *ROCK2* is shown. **F.** Fraction of migrated A549 cells for the indicated samples is shown as mean ± std from at least three independent experiments. One-way ANOVA, ****, *P*<0.0001, **, *P*<0.01, *, *P*<0.05, n.s., not significant. **G.** Fraction of migrated A549 cells (parental) and the derived single cell clones with the indicated FXR1 genotypes. Shown and quantified as in (**F**). One-way ANOVA, ****, *P*<0.0001, ***, *P*<0.001, *, *P*<0.05, n.s., not significant. The migration capacity of the single cell clones with WT genotype is significantly different from the parental cells. The migration capacity of the single cell clones with mutant FXR1 is significantly different from the WT clones. **H.** Western blot of the indicated endogenous proteins of the RhoA signaling pathway in A549 cells, grown in steady-state conditions and expressing the indicated shRNAs. α-Tubulin was used as loading control. **I.** Western blot of the indicated endogenous proteins of the RhoA signaling pathway in A549 cells after serum starvation and stimulation with thrombin for 10 minutes and expressing the indicated shRNAs. GAPDH was used as loading control. **J.** As in (**I**), but for shown for additional shRNAs. RLC T19 phosphorylation requires the presence of ROCK1, ROCK2, and FXR1, whereas FXR1 KD did not change MYPT1 T853 phosphorylation level. **K.** Active RhoA (RhoA-GTP) pulldown assay was performed in A549 cells expressing the indicated shRNAs, which were serum-starved and treated with LPA for 5 minutes. The level of active RhoA after GPCR activation is FXR1-independent. **L.** Validation of the indicated RLC and ROCK2 antibodies for PLA assay using immunofluorescence staining in A549 cells expressing the indicated shRNAs. The dilution factor used for each antibody is shown. Scale bar, 40 µm.

**Figure S7.**
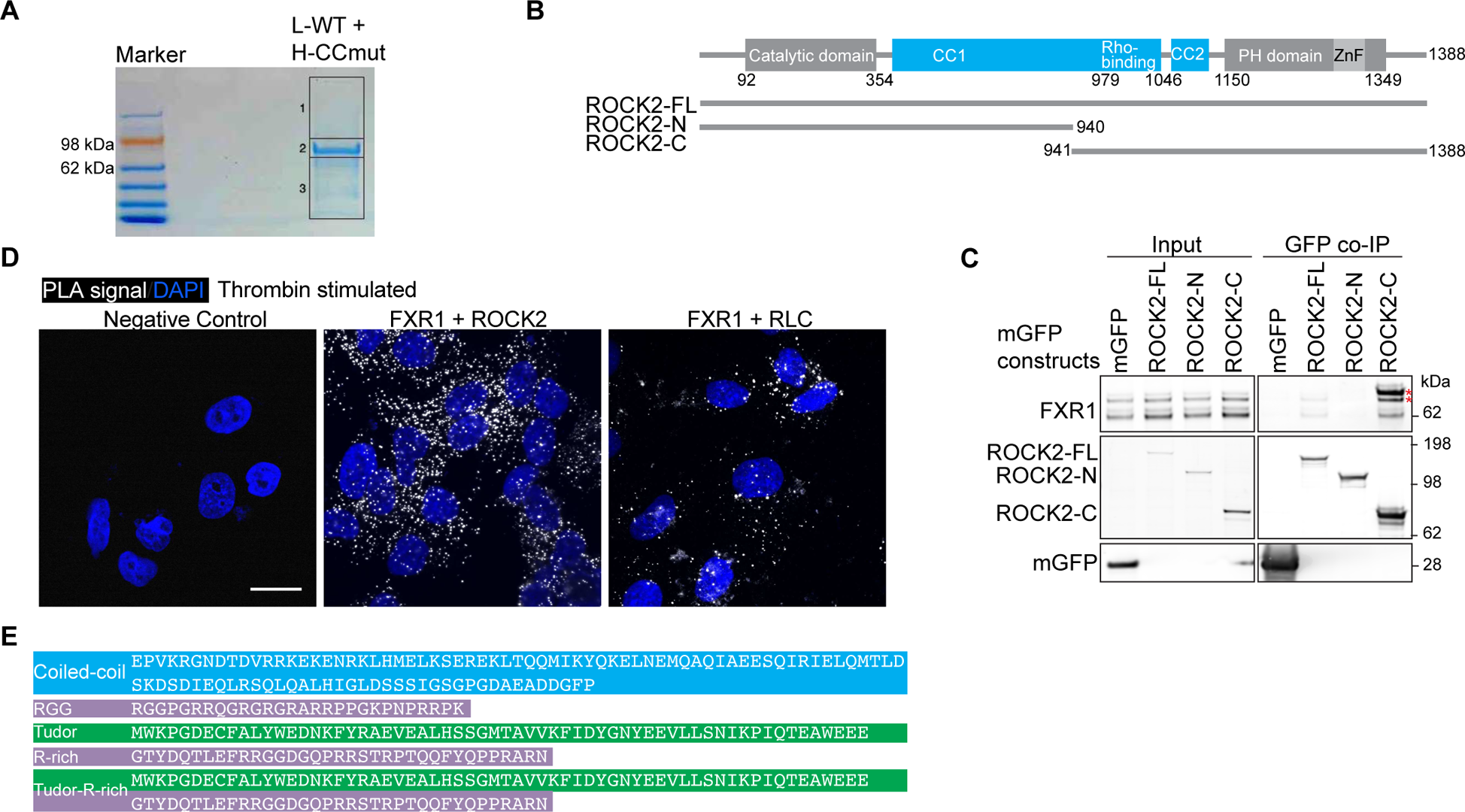
Proteins with binding sites for FXR1 are recruited into the FXR1 network, related to Figure 7. **A.** Coomassie staining of the gel used for SILAC proteomics prepared from HeLa cells. The three boxed areas represent the three gel slices processed for mass spectrometry analysis. **B.** Schematic of ROCK2 protein domains and GFP-ROCK2 constructs used. **C.** GFP co-IP of endogenous FXR1 protein after ectopic expression of GFP or the GFP-tagged ROCK2 constructs from (**B**) in HeLa cells. The two red star symbols mark a bleed-through signal from the blot for ROCK2-C. **D.** PLA performed in serum-starved and thrombin-stimulated A549 cells, indicating proximity between FXR1 and RLC as well as FXR1 and MYPT1. As negative control, the FXR1 antibody alone was used. DAPI staining visualizes the nucleus. Representative images of three independent experiments are shown. Scale bar, 20 µm. **E.** The amino acid sequences of the CC, Tudor, RGG, R-rich, and Tudor-R-rich domains fused to the C-terminus of GAPDH are shown. This panel is related to Fig. 7**D-F**.

## STAR methods

### RESOURCE AVAILABILITY

#### Lead contact

Further information and requests for resources and reagents should be directed to and will be fulfilled by the Lead contact, Christine Mayr.

#### Materials availability

- Plasmids generated in this study will be deposited to Addgene.
- Plasmids generated in this study not available at Addgene are available from the Lead Contact.
- The FXR1 knockin cell lines and the FXR1-N202S and FXR1-G266E cell lines (together with the control cell lines) generated in this study are available from the Lead Contact with a completed Materials Transfer Agreement.

#### Data and code availability

- The data of the TMT mass spectrometry experiment were deposited in the MassIVE repository (dataset identifier MSV000093384). The data of the SILAC mass spectrometry experiment were deposited in the MassIVE repository (dataset identifier MSV000093385).
- The HeLa RNA-seq sample and the FXR1 iCLIP data obtained from HeLa cells are available at ArrayExpress (ArrayExpress accession E-MTAB-13545).
- Western blot data, raw imaging data and scripts for analysis will be deposited at Mendeley.
- Any additional information required to reanalyze the data reported in this paper is available from the lead contact upon request.

### EXPERIMENTAL MODEL AND STUDY PARTICIPANT DETAILS

#### Cell lines

HeLa, a human cervical cancer cell line of female origin, was a gift from the Jonathan S. Weissman lab (UCSF), provided by Calvin H. Jan. HEK293T, a human immortalized embryonic kidney cell line of female origin, was purchased from ATCC. A549, a human lung cancer cell line of male origin, and MCF7, a human breast cancer cell line, were gifts from the lab of Robert Weinberg (Whitehead Institute). U2OS and U2OS FXR1, FXR2, and FMR1 triple knockout (U2OS ΔΔΔ) cell lines were a gift from the lab of Shawn Lyons (Boston University)^24^. All above cell lines were maintained at 37°C with 5% CO2 in Dulbecco’s Modified Eagle Medium (DMEM) containing 4,500 mg/L glucose, 10% heat inactivated fetal bovine serum, 100 U/ml penicillin and 100 μg/ml streptomycin. The human lung squamous cell lines EBC-1 and HCC95 were gifts from the Anti-tumor Assessment Core Facility and the lab of Charles Rudin (MSKCC). They were maintained in RPM1-1640 medium containing 10% heat-inactivated fetal bovine serum, 100 U/ml penicillin, and 100 μg/ml streptomycin. These cell lines have not been authenticated. The human iPSC cell line 731.2B was obtained from the SKI Stem Cell Research Facility at MSKCC^67^. The cells were maintained at 37°C with 5% CO2 in Stemflex medium (Thermo Fisher, A3349401). All cell culture vessels were coated with hESC-qualified Matrigel (Fisher Scientific, 354277). ROCK inhibitor (Y-27632, 10 μM, Stemcell Technologies, 73202) was added to the medium when the cells were passaged with 0.5 mM EDTA.

#### Constructs

##### GFP fusion constructs

All GFP fusion constructs were generated in the pcDNA3.1-puro-EGFP backbone as N-terminal fusion proteins with the original AUG omitted^7^. Monomeric (mGFP) was generated through the A207K mutation in EGFP and used in all constructs.

Human FXR1 mRNA was PCR-amplified from a HEK293T cDNA library and inserted between BsrGI and XhoI sites. The cDNA library was created with qScript cDNA SuperMix (Quantabio, 95048). A total of three isoforms were identified through Sanger sequencing: FXR1 isoform a (NM_005087.3, 621 amino acids (aa)), isoform b (NM_001013438.3, 539 aa), and isoform X4 (XM_005247816.3). If not stated otherwise, FXR1 isoform a was used. The FMR1 isoform 1 (NM_002024.6, 632 aa) coding sequence was amplified from the plasmid #48690 (Addgene) and inserted between BsrGI and EcoRV sites.

The GAPDH, TOP3B, and TDRD3 coding sequences were amplified from a HeLa cDNA library and inserted into the pcDNA3.1-puro-EGFP vector. The N-terminus of ROCK2 (aa 1-940) was amplified from the plasmid #70569 (Addgene) and cloned into the XhoI-linearized backbone with Gibson assembly master mix (E2621L, NEB) to obtain pcDNA3.1-puro-mGFP-ROCK2-N. The C-terminus of ROCK2 (aa 941-1388) was amplified from an A549 cDNA library and inserted between BsrGI and EcoRV sites. These two libraries were created by SuperScript IV VILO First-Strand Synthesis System (Invitrogen, 11756050). The N-terminus of ROCK2 was also amplified, and Gibson assembled into BsrGI-linearized pcDNA3.1-puro-mGFP-ROCK2-C to obtain the full-length ROCK2 construct. To generate pcDNA3.1-puro-mGFP-ROCK2-C-ΔCC, a gene fragment derived from the sequence between the SpeI and BbvCI sites of ROCK2-C, which lacked the sequence of the coiled-coil domain (aa 1046-1150) was synthesized (Genewiz). The exact sequence is listed in Table S4. This fragment and the pcDNA3.1-puro-ROCK2-C backbone were digested with SpeI and BbvCI. Since the backbone contained two SpeI sites, two of the three resulting fragments were collected, and the 490 bp fragment between SpeI and BbvCI was discarded. The other two fragments and the synthesized fragment were then ligated.

The FXR1 and FMR1 N- and C-terminal truncation constructs as well as the CC mutants were generated using PCR amplification of the desired coding sequence fragments and were subcloned into the pcDNA3.1-puro-mGFP backbone. Single point mutations to prolines in coiled-coil domains were introduced at the first amino acid of the predicted heptads. The exact mutated residues are detailed in Fig. S4B and in the list of plasmids in the Key Resource Table. Specific point mutations and coiled-coil swapping constructs were generated using pcDNA3.1-puro-mGFP-FXR1a or FMR1 via site-directed mutagenesis with Pfu Ultra HF DNA polymerase (Agilent). The second coiled-coil domain in FXR1 contains highly conserved residues in the first predicted heptad (Fig. S4A). This heptad was not disturbed when generating the CC mutants. The amino acid sequences of all FXR1 CC mutants are detailed in Fig. S4B.

GAPDH-fusion protein constructs were generated using Gibson assembly master mix with EcoRI linearized pcDNA3.1-puro-EGFP-GAPDH and desired PCR-amplified fragments. The amino acid sequences appended to GAPDH are shown in Fig. S7E. The pcDNA3.1-UBAP2L-mGFP construct was a gift from Christopher Hammell (CSHL). All constructs were verified by Sanger sequencing or whole plasmid sequencing. All oligos used for cloning are listed in Table S4.

##### shRNA constructs

A control shRNA against luciferase (MISSION® shRNA SHC007) was purchased. All other shRNAs were designed with the Broad Institute GPP web portal. DNA oligonucleotides listed in Table S4 were used as shRNA precursors and inserted into a backbone pLKO.1 vector (TRCN0000160812) between SgrAI and EcoRI sites. All vectors were verified by Sanger sequencing with U6 primer.

### Transfection

Besides CRISPR-based gene editing experiments, all transfections into HeLa and U2OS ΔΔΔ cells were performed with Lipofectamine 2000 (Invitrogen, 11668019).

For testing the amount of pcDNA3.1-mGFP-FXR1a plasmid to transfect to mimic endogenous FXR1 level, 500, 250, 125, and 62.5 ng of plasmid was mixed with 3 µl Lipofectamine, respectively, and transfected into HeLa grown in 35 mm dishes. 250 ng was determined to be the optimal amount. For all imaging-related experiments, 50 ng of FXR1 plasmid was mixed with 0.6 µl Lipofectamine to transfect one well of a 24-well plate. For other experiments, amounts were scaled up according to the surface area of the dish. For pcDNA3.1-puro-mGFP-FMR1, 100 ng plasmid per well of a 24-well plate was transfected.

For GFP trap mediated co-immunoprecipitation, 6 µg DNA of TDRD3, UBAP2L, TOP3B, GAPDH, or GAPDH-fusion constructs was transfected into HeLa cells seeded in 10 cm dishes with 10 µl Lipofectamine 2000 in a total of 1 ml OPTI-MEM (Gibco, 31985062).

### shRNA-mediated knockdown

Stable cell lines were generated for shRNA-mediated knockdown experiments. 2 µg pLKO.1 plasmid was co-transfected with 1.8 µg pCMV-dR8.2 and 0.2 µg pCMV-VSV-G with 7 µl Lipofectamine 2000 into HEK293T cells seeded in 6-well plates one day ahead. The medium was changed 6 hours after transfection. Viral particles were harvested 48 hours after transfection by filtering through a 0.45 µm filter unit. 50 to 100 µl viral particles were used to transduce target cells grown in 6-well plates in the presence of 8 µg/ml polybrene. 24 hours after transduction, puromycin was added to the medium at a final concentration of 2 µg/ml for HeLa and A549 cells to select for shRNA-expressing cells. Cells were expanded into media containing 1 µg/ml of puromycin for maintenance after two days of selection.

### siRNA-mediated knockdown

All siRNAs were ordered from Sigma-Aldrich, either predesigned or customized. MISSON siRNA Universal Negative Control #1 (SIGMA, SC001) was used as a negative control. The sequences of the used siRNAs are listed in Table S4. siRNAs were transfected with Lipofectamine RNAiMAX (Invitrogen, 13778150) at a final concentration of 15 nM following the manufacturer’s instructions. Cells were harvested three days after transfection for Western blotting or live cell imaging.

### CRISPR-Cas9-mediated knockin of GFP or NG

#### mGFP-FXR1 or mNG-FXR1 knockin cells

Three gRNAs were designed with CRISPOR and ordered from IDT^68^. All three gRNAs worked efficiently and generated mGFP-FXR1 expressing cells with an indistinguishable microscopic distribution of the endogenous fusion protein. All reported knockin cell lines in this work were generated with sgRNA1 (Table S4). The repair donor gBLOCK was designed to include the desired tag (mGFP or mNeonGreen) with a 500 bp overhang on each side for homologous recombination. The donor sequences are listed in Table S4). Silent mutations disrupting the PAM sequences of all three gRNAs were introduced. The gBLOCK was synthesized at Genewiz and cloned into pUC-GW-AMP. The final double-stranded DNA donor was produced using PCR amplification with Q5 HF DNA polymerase (NEB, M0491) and the forward and reverse oligos (KI-donor-F and KI-donor-R) (Table S4).

For transfection, cells were seeded in 12-well plates one day ahead. 1.25 µg Cas9 protein (IDT #1078728) and 315 ng sgRNA (IDT synthesized) were mixed with 125 µl Opti-MEM for 10 minutes (min). Up to 2.5 µg dsDNA donor and 4 µl TranxIT X2 transfection reagent (Mirus, MIR6003) were added to the mixture, incubated for 15 min at room temperature, and added to HeLa, HEK293T, or A549 cells. Transfected cells were submitted to FACS sorting at least five days after transfection to collect mGFP- or mNeonGreen-positive cells. GFP-positive bulk cells were used. Successful knockin was confirmed with confocal microscopy, western blotting, and genotyping, followed by sequencing. The primers used for genotyping are listed in Table S4.

### Mutation of endogenous FXR1 using base editing

To disrupt the first coiled-coil domain of human FXR1 in A549 cells, base editing was used to change N202 to S202. Adenine Base Editor ABEmax(7.10)-SpG-P2A-EGFP was expressed from the Addgene plasmid #140002^69^. FXR1 exon 7 specific sgRNAs were designed with CRISPOR^68^ and expressed from the backbone BPK1520 (Addgene #65777) driven by the U6 promoter. DNA oligos used for cloning are listed in Table S4. gRNAs were annealed and phosphorylated, then ligated into BsmBI-digested and dephosphorylated BPK1520 backbone.

Transfections were performed between 20 and 24 hours after seeding 4 x 10^5^ HEK293T or A549 cells in 6-well plates. 1.4 µg of base-editor and 600 ng of sgRNA expression plasmids were mixed with 15 µl of TransIT-X2 (Mirus, MIR6003) in a total volume of 300 µl Opti-MEM, incubated for 15 min at room temperature and added to A549 cells. Transfected cells were submitted to FACS sorting five days after transfection to collect GFP-positive cells. To perform FACS sorting, cells in 10 cm dishes were washed with PBS and trypsinized with 2 ml trypsin at room temperature for 5 min. After carefully removing trypsin, the cells were resuspended in 2 ml FACS buffer (growth media containing 2.5% FBS) and passed through a cell strainer. GFP-positive cells were sorted in bulk and 96-well plates with one cell per well on a BD FACSymphony^TM^ S6 cell sorter.

To assess base editing efficiency, one week after sorting, genomic DNA was extracted using QuickExtract DNA extraction solution (LGC, SS000035-D2) from the bulk sorted cells. CRISPRseq DNA was PCR amplified with Q5 (NEB) using oligos listed in Table S4, ran on an agarose gel, and gel purified using QIAquick Gel Extraction Kit. CRISPRseq results were processed using the CRISPRESSO2 pipeline^70^. Single cell-derived clones were obtained through FACS sorting, expanded, and genotyped with Sanger sequencing. For Sanger sequencing, the forward oligo for amplicon generation was used as the sequencing primer. Two WT control *FXR1* clonal cell lines, three heterozygous *FXR1-WT/FXR1-N202S*, and one homozygous *FXR1-N202S/FXR1-N202S* cell line were generated and used in this study.

### Mutation of endogenous FXR1 using prime editing

Prime editing was employed to install the mutation FXR1-G266E at the endogenous locus in A549 cells. Prime editor PEmax with P2A-EGFP was expressed from the addgene plasmid #180020. The epegRNA was designed with PE-designer^71^ and expressed from the backbone pU6-tevopreq1-GG-acceptor (Addgene # 174038) driven by the U6 promotor. The extra nicking gRNA was expressed from the backbone LsgRNA (Addgene #47108).

Transfections were performed between 20 and 24 hours after seeding 4 x 10^5^ cells in 6-well plates. 4 µg of prime editor, 1.3 µg of epegRNA, and 440 ng of nicking gRNA expression plasmids were mixed with 10 µl of TransIT-X2 (Mirus, MIR6003) in a total volume of 300 µl Opti-MEM, incubated for 15 min at room temperature and added to A549 cells. Transfected cells were submitted to FACS sorting five days after transfection to collect GFP-positive cells. GFP-positive cells were sorted in 96-well plates with one cell per well on a BD FACSymphony^TM^ S6 cell sorter.

To genotype the resulting single cell-derived clones, amplicons were generated with oligos FXR1-KH1-F and FXR1-KH1-R (listed in Table S4). The PCR products were sequenced using the oligo FXR1-KH1-F with Sanger sequencing. Three WT control FXR1 clonal cell lines and two heterozygous *FXR1-WT/FXR1-G266E* cell lines were generated and used in this study.

### Immunofluorescence staining

Cells were seeded in 4-well chamber slides (Millipore, PEZGS0416). Specifically, for HEK293T cells, the chambers were coated with 0.01% Poly-L-lysine (Sigma, P4707) at room temperature for one hour before seeding. The day after, cells were washed in PBS (-Ca^2+^, -Mg^2+^), fixed in 4% PFA for 10 min at room temperature, and washed twice with PBS. The cells were then permeabilized in 0.1% Triton X-100 in PBS for 7 min. After washing three times with PBST (PBS with 0.1% Tween-20), the cells were incubated in the blocking buffer (3% BSA in PBST) for 1 hour. The cells were then incubated in primary antibody diluted in the blocking buffer for 3 hours at room temperature or overnight at 4°C. After washing the cells three times in PBST, the cells were incubated with secondary antibody diluted at 1:1000 in blocking buffer for 1 hour. The cells were washed three times with PBST and mounted in ProLong Gold Antifade Mountant with DAPI (Invitrogen, P36941) with precision cover glasses No. 1.5H (Marienfeld, 0107222). All antibodies are listed in the Key Resource Table.

### Confocal microscopy

Two confocal microscopes were used depending on the availability. Most live cell imaging was conducted on the ZEISS LSM880 confocal laser scanning microscope in Airyscan mode at 37°C with a Plan-Apochromat 63x/1.4 Oil objective (Zeiss), driven by ZEN black. Exceptions are data shown in Fig. 3I and Supplemental Videos S1-S6, which were acquired with a SoRa spinning disk microscope. Most fixed samples were imaged with a SoRa spinning disk microscope. The SoRa spinning disk was equipped with an ORCA-Fusion BT Digital CMOS camera (C15440-20UP, Hamamatsu), a motorized piezo stage, and 63x/1.40 CFI Plan Apo oil immersion objective, driven by the software NIS-ELEMENTS (Nikon).

For live cell imaging with LSM880, including FRAP experiments, cells were seeded in 4-well Nunc Lab-Tek II chambered coverglasses (Thermo Scientific, 155360) and transfected with constructs with the above-mentioned amount. Fourteen to 17 hours after transfection, cells were mounted on the stage housed in a live cell imaging chamber (Zeiss) at 37°C and 5% CO2. Z stack images were captured with an interval size of 160 nm when applicable. Excitations were performed sequentially using 405, 488, 594, or 633 nm laser, and imaging conditions were experimentally optimized to minimize bleed-through. For live cell imaging with SoRa, the cells were seeded in Ibidi µ-Slide 4-well chambers (Ibidi USA, NC0685967) using FluoroBrite™ DMEM (Gibco, A1896701). The samples were excited with the 488 nm laser and exposed for 80 ms. Raw images are presented unless otherwise stated.

#### Imaging after RNase A treatment

The cells were seeded in a glass-bottomed 4-well chamber and transfected with 50 ng mGFP-FXR1-N2 construct. 15 hours after transfection, the cells were washed twice with PBS, then washed once more with “transport buffer”, which contains 20 mM HEPES pH 7.4, 100 mM potassium acetate, 3.5 mM magnesium acetate, 1 mM EGTA, and 250 mM sucrose. The cells were permeabilized with 500 µl of the above-mentioned buffer containing 50 µg/ml digitonin for 1 min. The cells were washed twice with PBS and incubated in PBS supplemented with or without 1 mg/ml RNase A (Sigma-Aldrich, Cat# R4642). The signal obtained from the GFP-FXR1-N2 construct was recorded with the ZEISS LSM880 confocal laser scanning microscope. At 30 min post RNase A addition, the assembled network was fully dissociated into spherical granules.

#### Fluorescence recovery after photobleaching (FRAP)

HeLa cells were seeded in 4-well Nunc Lab-Tek II chambered coverglass (Thermo Scientific, 155360). FRAP experiments were performed with ZEISS LSM880 in the airyscan mode using the 488 nm laser. A square area of 0.5 x 0.5 µm^2^ was bleached with maximal power. For full-length FXR1, the bleaching area was 1.6 x 1.6 µm^2^. The fluorescence signal was acquired at the maximum speed possible for 100 seconds at an interval of 2 seconds. The fluorescence intensity of the bleached area was extracted with ZEN software black edition (ZEISS). The prebleached fluorescence intensity was normalized to one, and the signal after bleach was normalized to the pre-bleach level. No photobleaching was observed on non-bleached areas; we therefore took the Plateau values as mobile fractions.

#### Three-dimensional colocalization

FMR1 and FXR1 were stained in HeLa cells, and stacks of images were acquired with a step size of 0.2 µm on a SoRa spinning disk microscope. A 63x/1.40 CFI Plan Apo oil immersion objective and a 4x magnification changer for SoRa were used. Images were deconvolved with default settings using NIS-elements software. These images were then imported into Imaris software and automatically thresholded. The ‘3D coloc’ function was applied to all volumes and generated related parameters, including the percentage of volume colocalized for FMR1 and FXR1, as well as the Person’s correlation coefficient in the thresholded volume.

#### Connected component analysis

Confocal images of GFP-FXR1-N1 or -N2 were acquired with either LSM880 or SoRa. The images were then analyzed in Python with scikit-image^72^. Briefly, the images were automatically thresholded and the connected components, which are called objects in this paper, were identified using the ‘skimage.measure’ function with the connectivity specified as 2. These objects were then assigned random colors. Each object’s size (area) and the total number of objects per cell were extracted.

### RhoA pathway stimulation and stress fiber staining

A549 cells were seeded in 4-well chamber slides (Millipore, PEZGS0416) at a density of 0.03 x10^6^ cells per well. The evening after, the cells were washed twice with starvation media (DMEM-HG without FBS) and incubated in 500 µl starvation media for 17 hours. The cells were stimulated with 3 µM LPA (Avanti, 857130P) or 60 nM thrombin (Novagen, 69671). 30 min after stimulation, the cells were washed with PBS and fixed in 4% PFA for 10 min at room temperature. Filamentous actin was stained with Phalloidin-iFluor 555 Reagent (Abcam, ab176756) per the manufacturer’s instructions. The presence of stress fibers for each cell was scored either positive or negative. A fraction of the dataset was blindly scored by two authors, and a similar fraction of stress fiber-positive cells was found. Most of the images were scored by the first author.

For western blot analysis, cells were seeded in 6-well plates lysed in 1x reducing Laemmli SDS sample buffer 15 min after stimulation unless otherwise stated.

### Active RhoA pulldown

Active RhoA pulldown was performed with the RhoA Pull-Down Activation Assay Kit (Cytoskeleton, Inc, BK036-S), following the manufacturer’s instructions. Briefly, A549 cells were seeded in 6 cm dishes and serum-starved for 17 hours. The cells were then stimulated with or without 3 µM LPA for 5 min before washing and lysing. Active RhoA was enriched by GST-tagged Rhotekin-RBD protein coupled to agarose beads. The beads were thoroughly washed, and the resulting products were separated on SDS-PAGE and analyzed using Western blotting.

### Proximity ligation assay

A549 cells were seeded onto glass coverslips (Fisherbrand, 12541001, No 1.5) with a 12 mm diameter placed in 24-well plates. The cells were serum-starved for 17 hours before stimulation. 10 min after 60 nM thrombin stimulation, cells were fixed with 4% PFA in PBS for 10 min, permeabilized with 0.1% Triton in PBS for 7 min, washed with PBST three times, blocked in 3% BSA in PBS for 30 min, and incubated with primary antibody diluted in blocking buffer overnight at 4°C. The next day, cells were washed with PBST three times and incubated with secondary antibody with PLUS and MINUS DNA probes (Sigma-Aldrich, DUO92102) for 1 hour at 37°C. Washed with Wash Buffer A two times, incubated in ligation mix for 30 min at 37°C. Washed with Wash Buffer A two times, incubated in signal amplification mix for 100 min at 37°C. Finally, washed with Wash Buffer B two times, and with 0.01 x Wash Buffer B once. Cells were then mounted in Prolong Gold Antifade Mountant with DAPI for imaging on a confocal microscope. Z-section images (*N* = 21) separated by 0.4 µm increments were captured. Images were analyzed in ImageJ with a custom script. Briefly, images were max-z projected and auto-thresholded. The dots were then selected with the ‘find maxima’ function and counted for individual cells with manually drawn regions of interest (ROIs) using the ROI manager.

### Migration assay

6,000 serum-starved A549 cells in 100 µl serum-free DMEM were dispensed into the transwell insert in a 24-well plate (Costar 3422, 8 µm pore size) with 500 µl complete DMEM. When used, ROCK inhibitor Y-27632 (Catalog # 72307, STEMCELL technologies) was added at a final concentration of 10 µM for two hours prior to dispensing into transwell inserts. And fresh ROCK inhibitor was added to the transwells for the whole duration of the experiment. The wells, and the inserts were washed with PBS 20 hours after seeding. 500 µl accutase (Innovative Cell Technologies, AT-104) was added to the wells and incubated at room temperature for 8 minutes. Cells in the accutase solution were collected by centrifugation and subjected to Cyquant (Invitrogen, C7026) based DNA quantity measurement using a plate reader SpectraMax iD5 and clear bottom black assay plates (Costar, 3603).

### Size exclusion chromatography

8 x 10^6^ HeLa cells or A549 cells were lysed in 550 µl mild lysis buffer containing 50 mM HEPES, pH 7.4, 150 mM NaCl, 0.5% NP40, 1 mM PMSF, and 1 x EDTA-free Protease Inhibitor Cocktail (Roche). The cells were further broken down with six passes through a 27-gauge needle. The lysate was cleared at top speed for 10 min with a tabletop centrifuge at 4°C. 500 µl crude lysate was loaded into the Superose^®^ 6 Increase 10/300 GL column (Cytiva, 29091596) driven by an AKTA FPLC system (GE Healthcare). 1 ml fractions were collected over the entire run. 200 µl 100% (w/v) TCA (SIGMA, T9159) was added to each fraction and kept at −80°C overnight. The precipitated protein was collected and washed twice with 1 ml of ice-cold acetone. Finally, protein was airdried and resuspended in 120 µl 2x reducing Laemmli SDS sample buffer. These samples were further analyzed using western blotting.

When fractionating GFP-FXR1 WT and CC mutant fusion proteins, instead of TCA precipitation, 150 µl of each collected fraction was loaded into a 96-well solid black microplate (Corning, 3915) and analyzed with an Infinite M1000 plate reader (Tecan). Fluorescence was collected with top reading mode, excited at 488 nm, and collected at 510 ± 5 nm with optimal gain. A GFP negative lysate sample from the same cell type was fractionated and served as background control for the autofluorescence.

### Co-immunoprecipitation

GFP trap (Chromotek, Gta-100) co-IP was performed as follows. HeLa cells were transfected with constructs expressing GFP or GFP-fusion proteins, as described above. About 17 hours after transfection, the cells were washed with PBS twice and drained of the remaining liquid. The cells were scrapped into 700 µl lysis buffer containing 50 mM HEPES, pH 7.4, 150 mM NaCl, 1% NP40, 0.5% sodium deoxycholate, 0.05% SDS, 1 mM EDTA, 1 x EDTA-free protease inhibitor cocktail (Roche). The cells were lysed on ice for 30 min. After centrifugation at 21,130 g for 10 min, GFP-trap co-IP was performed following the manufacturer’s instructions with 15 µl slurry per reaction. GFP-trap beads were added, incubated with cell lysate for 1 to 2 hours at 4°C on a rotator, and washed four times with ice-cold wash buffer containing 50 mM HEPES, pH 7.4, 150 mM NaCl, and 1 mM EDTA.

When RNase A treatment was required, the beads were split into two samples after the third wash and resuspended in 200 µl of wash buffer. A final concentration of 30 mg/ml of RNase A (Sigma-Aldrich, Cat# R4642) was added and treated at room temperature for 30 min. After a final wash, the GFP-trap beads were mixed with 2x Laemmli sample buffer, boiled at 95°C for 5 min, and subjected to Western blotting.

### Western blotting

Cells were washed with PBS and lysed with 1x reducing Laemmli SDS sample buffer (Thermo Scientific Chemicals, J60015-AC) to generate whole cell lysate. The viscous products were transferred to Eppendorf tubes and boiled at 95°C for 15 min.

For co-immunoprecipitation experiments, proteins were eluted from beads by boiling in 2x reducing Laemmli SDS sample buffer at 95°C for 5 min.

Denatured protein samples were separated in 4%-12% NuPAGE Bis-Tris gels (Invitrogen) and wet-transferred to nitrocellulose membranes with X cell II blot module (Invitrogen). For analyzing high molecular weight proteins such as Myosin (MYH9), samples were separated in 3% - 8% Tris-Acetate gels (Invitrogen) with NuPAGE Tris-Acetate SDS Running buffer. Membranes were blocked with Odyssey blocking buffer (LI-COR) or 5% non-fat milk in TBST (exclusively when blotting RLC and pRLC) and then incubated with primary antibody at 4°C overnight. Membranes were washed three times with PBST (0.1% Tween) and incubated with dye-labeled secondary antibody. Membranes were scanned with the Odyssey DLx system (LI-COR). All antibodies are listed in the Key Resource Table.

### Oligo(dT) pulldown of mRNA-associated proteins without cross-linking

Plasmids expressing GFP-fusion proteins were transfected into U2OS *FXR1/FXR2/FMR1* triple knockout cells one day ahead. About 6 x 10^6^ U2OS were harvested for each reaction. For A549 cells with endogenous FXR1-N202S or FXR1-G266E mutation, the cells were seeded one day ahead to reach 70% confluency the next day for harvesting. Cells were washed with ice-cold PBS and lysed in 0.7 ml ice-cold lysis buffer containing 50 mM HEPES, pH 7.4, 150 mM NaCl, 1% NP40, 0.5% sodium deoxycholate, 0.05% SDS, 1 mM EDTA, and 1 x protease inhibitor cocktail (Roche). Samples were further lysed with forty strokes of a chilled dounce homogenizer. Lysates were cleared at 21,000 x g for 10 min at 4°C. 30 µl oligo(dT)25 magnetic beads (NEB, S1419S) were equilibrated in wash buffer containing 50 mM HEPES, pH 7.4, 150 mM NaCl, and 1 mM EDTA. The cleared lysate was mixed with the beads and was rotated for 60 min at 4°C. Samples were washed four times with 0.7 ml wash buffer and eluted from the beads with 2x reducing Laemmili sample buffer at 95°C for 5 min. The samples were analyzed using western blotting.

### RNA immunoprecipitation without cross-linking

RNA immunoprecipitation assays were used to validate iCLIP results. 8 x 10^6^ HeLa cells per condition were homogenized in lysis buffer containing 50 mM HEPES, pH 7.4, 150 mM NaCl, 1% NP40, 0.5% sodium deoxycholate, 0.05% SDS, 1 mM EDTA, 1 x EDTA-free protease inhibitor cocktail (Roche), and 2 U/ml SUPERase•In™ RNase Inhibitor (Invitrogen). Cleared lysates were incubated with 10 µg anti-FXR1 antibody (Novus #NBP2-22246) or Rabbit IgG (Cell Signaling Technologies #2729)-coupled protein A beads for four hours at 4°C. After washing the beads three times with wash buffer containing 50 mM HEPES pH 7.4, 150 mM NaCl, and 0.05% NP40, RNA was eluted from the beads with 1 mg/ml proteinase K (AM2546) at 50°C for 40 min. RNA was then isolated with TRI reagent (Invitrogen) with standard procedure and reverse transcribed using qScript cDNA SuperMix (Quantabio). The primer sequences for RT-qPCR analysis are listed in Table S4. Enrichment relative to input RNA was calculated using cycle threshold values for each mRNA. The final fold change of FXR1/IgG was obtained by dividing the enrichment over input of FXR1-IP by IgG-IP.

### SILAC mass spectrometry

HeLa cells stably expressing shRNAs against FXR1 were cultivated in DMEM medium (Thermo Scientific, A33822) supplemented with 10% dialyzed FBS (Gibco, 26400044) and 1% penicillin and containing either “light” (L-Arginine-HCL (Thermo Scientific, 89989), L-Lysine-2HCL (Thermo Scientific, 89987)) or “heavy” (L-Arginine-HCL (13C6, 99%; 15N4, 99%; Cambridge Isotope Laboratories, CNLM-539-H-0.05, L-Lysine-2HCL (13C6, 99%; Thermo Scientific, 1860969)) stable isotope labeled amino acids. Cells were cultivated for at least six passages before the incorporation efficiency was verified by mass spectrometry analysis to be above 99%.

The ′light′ HeLa cells were transfected with GFP-FXR1a (containing a silent mutation that makes it shRNA-resistant), and the ′heavy′ cells were transfected with shRNA-resistant GFP-FXR1a-CC2 mutant (V361P) using Lipofectamine 2000. After 18 hours, transfected cells were collected and lysed in buffer containing 50 mM Tris–HCl pH 7.5, 150 mM NaCl, 1% Triton X-100, 1 mM EDTA, 0.25% sodium deoxycholate, and 1x protease inhibitor cocktail (Roche, 11836153001). GFP-trap (Chromotek, Gta-100) co-IP was performed separately using light and heavy lysates. 30 µl slurry per sample was used. The resulting beads were pooled and mixed with 2x Laemmli sample buffer followed by SDS-gel electrophoresis in MES running buffer using 4-12% Bis-Tris NuPAGE gels at 120 V for 10 min. Following the manufacturer’s instructions, the protein gels were stained with SimplyBlue (Life Technologies) and submitted to the MSKCC Proteomics Core facility for SILAC mass spectrometry analysis.

The samples were divided into three gel slices (Fig. S7A), and all three gel slices were processed for MS analysis. They were washed with 1:1 (Acetonitrile:100 mM ammonium bicarbonate) for 30 min, dehydrated with 100% acetonitrile for 10 min, excess acetonitrile was removed, and slices were dried in speed-vac for 10 min without heat. Gel slices were reduced with 5 mM DTT for 30 min at 56°C in a thermomixer (Eppendorf), chilled to room temperature, and alkylated with 11 mM IAA for 30 min in the dark. Gel slices were washed with 100 mM ammonium bicarbonate and 100% acetonitrile for 10 min each. Excess acetonitrile was removed and dried in speed-vac for 10 min without heat, and gel slices were rehydrated in a solution of 25 ng/μl trypsin in 50 mM ammonium bicarbonate on ice for 30 min. Digestions were performed overnight at 37°C in a thermomixer. Digested peptides were collected and further extracted from gel slices in an extraction buffer (1:2 (v/v) 5% formic acid/acetonitrile) at high-speed shaking in a thermomixer. Supernatant from both extractions was combined and dried in a vacuum centrifuge. Peptides were desalted with C18 resin-packed stage tips, lyophilized, and stored at −80°C until further use.

#### LC-MS/MS analysis

Desalted peptides were dissolved in 3% acetonitrile/0.1% formic acid and were injected onto a C18 capillary column on a nano ACQUITY UPLC system (Water), which was coupled to the Q Exactive plus mass spectrometer (Thermo Scientific). Peptides were eluted with a non-linear 200 min gradient of 2-35% buffer B (0.1% (v/v) formic acid, 100% acetonitrile) at a 300 nl/min flow rate. After each gradient, the column was washed with 90% buffer B for 5 min and re-equilibrated with 98% buffer A (0.1% formic acid, 100% HPLC-grade water). MS data were acquired with an automatic switch between a full scan and 10 data-dependent MS/MS scans (TopN method). The target value for the full scan MS spectra was 3 x 10^6^ ions in the 380-1800 *m/z* range with a maximum injection time of 30 ms and resolution of 70,000 at 200 *m/z* with data collected in profile mode. Precursors were selected using a 1.5 *m/z* isolation width. Precursors were fragmented by higher-energy C-trap dissociation (HCD) with a normalized collision energy of 27 eV. MS/MS scans were acquired at a resolution of 17,500 at 200 m/z with an ion target value of 5 x 10^4^, maximum injection time of 60 ms, dynamic exclusion for 15 s and data collected in centroid mode.

### Tandem Mass Tag (TMT) Multiplexed Quantitative Mass Spectrometry

The TMT analysis was performed with four replicates per sample. 4 x 10^6^ HeLa cells expressing control shRNA or an shRNA against FXR1 were used as samples. Cells were trypsinized and washed three times with ice-cold PBS. Pelleted cells were snap-frozen in liquid nitrogen after the final wash. Cell pellets were lysed with 200 μl buffer containing 8 M urea and 200 mM EPPS pH = 8.5, with protease inhibitor (Roche) and phosphatase inhibitor cocktails 2 and 3 (Sigma). Benzonase (Millipore) was added to a concentration of 50 µg/ml and incubated at room temperature for 15 min followed by water bath sonication. Samples were centrifuged at 4°C, 14,000 g for 10 min, and the supernatant was extracted. BCA assay (Pierce) was used to determine the protein concentration. Protein disulfide bonds were reduced with 5 mM tris (2-carboxyethyl) phosphine at room temperature for 15 min, then alkylated with 10 mM iodoacetamide at room temperature for 30 min in the dark. The reaction was quenched with 10 mM dithiothreitol, incubated at room temperature for 15 min. Aliquots of 100 µg were taken for each sample and diluted to approximately 100 μl with lysis buffer. Samples were subjected to chloroform/methanol precipitation as previously described^73^. Pellets were reconstituted in 200 mM EPPS buffer and digested with Lys-C (1:50 enzyme-to-protein ratio) and trypsin (1:50 enzyme-to-protein ratio) at 37°C overnight.

Peptides were TMT-labeled as described^73^. Briefly, peptides were TMT-tagged by adding anhydrous ACN and TMTPro reagents (16plex) for each respective sample and incubated for one hour at room temperature. A ratio check was performed by taking a 1 μl aliquot from each sample and desalted by StageTip method^74^. TMT tags were then quenched with hydroxylamine to a final concentration of 0.3% for 15 min at room temperature. Samples were pooled 1:1 based on the ratio check and vacuum-centrifuged to dryness. Dried peptides were reconstituted in 1 ml of 3% ACN/1% TFA, desalted using a 100 mg tC18 SepPak (Waters), and vacuum-centrifuged overnight.

Peptides were centrifuged to dryness and reconstituted in 1 ml of 1% ACN/25mM ABC. Peptides were fractionated into 48 fractions. An Ultimate 3000 HPLC (Dionex) coupled to an Ultimate 3000 Fraction Collector using a Waters XBridge BEH130 C18 column (3.5 µm 4.6 x 250 mm) was operated at 1 ml/min. Buffer A, B, and C consisted of 100% water, 100% ACN, and 25 mM ABC, respectively. The fractionation gradient operated as follows: 1% B to 5% B in 1 min, 5% B to 35% B in 61 min, 35% B to 60% B in 5 min, 60% B to 70% B in 3 min, 70% B to 1% B in 10min, with 10% C the entire gradient to maintain pH. The 48 fractions were then concatenated to 12 fractions, (i.e. fractions 1, 13, 25, 37 were pooled, followed by fractions 2, 14, 26, 38, etc.) so that every 12^th^ fraction was used to pool. Pooled fractions were vacuum-centrifuged and then reconstituted in 1% ACN/0.1% FA for LC-MS/MS.

Fractions were analyzed by LC-MS/MS using a NanoAcquity (Waters) with a 50 cm long (inner diameter 75 µm) EASY-Spray Column (PepMap RSLC, C18, 2 µm, 100 Å) heated to 60°C coupled to an Orbitrap Eclipse Tribrid Mass Spectrometer (Thermo Fisher Scientific). Peptides were separated by direct injection at a flow rate of 300 nl/min using a gradient of 5 to 30% acetonitrile (0.1% FA) in water (0.1% FA) over three hours and then to 50% ACN in 30 min and analyzed by SPS-MS3. MS1 scans were acquired over a m/z 375-1500 range, 120K resolution, AGC target (standard), and maximum IT of 50 ms. MS2 scans were acquired on MS1 scans of charge 2-7 using isolation of 0.5 m/z, collision-induced dissociation with activation of 32%, turbo scan, and max IT of 120 ms. MS3 scans were acquired using specific precursor selection (SPS) of 10 isolation notches, m/z range 110-1000, 50K resolution, AGC target (custom, 200%), HCD activation of 65%, max IT of 150 ms, and dynamic exclusion of 60 s.

### FXR1 iCLIP

HeLa cells were seeded in 10 cm dishes to reach 70% confluency the next day. After 24 hours, cells were transfected with either mGFP-FXR1-WT or mGFP-FXR1-CC2mut, as described in the transfections section.

20 hours after transfection, cells were rinsed once with ice-cold PBS and 6 ml of fresh PBS was added to each plate before proceeding to the improved iCLIP protocol^75^, with the following details. Cells were irradiated once with 150 mJ/cm^2^ in a Spectroline UV Crosslinker at 254 nm. Irradiated cells were scraped into Eppendorf tubes, spun at 500 x g for one minute, and snap-frozen in liquid nitrogen. Crosslinked cell pellets were lysed in iCLIP lysis buffer (50 mM Tris-HCl pH 7.4, 100 mM NaCl, 1% Igepal CA-630 (Sigma I8896), 0.1% SDS, 0.5% sodium deoxycholate, Complete protease inhibitor cocktail (Roche, 5056489001) and sonicated with the Bioruptor Pico for 10 cycles 30 seconds ON/30 seconds OFF. For RNA fragmentation, 4 U of Turbo DNase (Ambion, AM2238) and 0.1 U of RNase I (Thermo Scientific, EN0601) were added per 1 mg/ml lysate. Lysates were pre-cleared by centrifugation at 20,000 x g at 4°C. A mix of Protein G Dynabeads (100 µl per sample, Life Technologies) was coupled to 4 µg of rabbit anti-GFP antibody (Abcam ab290) and used for FXR1 protein-RNA complexes immunoprecipitation. Bead bound complexes were washed with high salt (50 mM Tris-HCl, pH 7.4, 1 M NaCl, 1 mM EDTA, 1% Igepal CA-630 (Sigma I8896), 0.1% SDS, 0.5% sodium deoxycholate) and PNK wash buffer (20 mM Tris-HCl, pH 7.4, 10 mM MgCl2, 0.2% Tween-20). RNA was first dephosphorylated and then ligated to a pre-adenylated infra-red labeled L3-IR adaptor on beads^76^. Excess adaptor was removed by incubation with 5′ deadenylase (NEB M0331S) and the exonuclease RecJf (NEB M0264S). GFP-FXR1 protein-RNA complexes were eluted from the beads by heating at 70°C for one minute, size-separated with SDS-PAGE, transferred to a nitrocellulose membrane, and visualized by iBright Imaging Systems via the infrared-labeled adaptor. RNA was released from the membrane by proteinase K digestion and recovered by precipitation. cDNA was synthesized with Superscript IV Reverse Transcriptase (Life Technologies) and circularized by CircLigase II. Circularized cDNA was purified with AMPure XP beads (A63880, Beckman Coulter), amplified by PCR, size-selected with AMPure beads, and quality-controlled for sequencing. Libraries were sequenced as single-end 100 bp reads on Illumina HiSeq 4000.

### Data analysis

#### Protein domains

##### Protein disorder prediction

Regions containing IDRs were determined using IUPred2A prediction program using ‘long disorder’^77^. Regions with scores higher than 0.5 are considered disordered.

##### Coiled-coil domain prediction

CC domains were predicted using the ‘coils’ program. A CC domain was predicted when the coils score was greater than 0.2. Proteins with predicted CC domains are listed in Table S1. Based on these criteria, 4168 (out of 8901 expressed genes) in HeLa cells encode proteins with at least one predicted CC domain. The resulting expected frequency of CC domains is 0.468. As this prediction program is no longer available, CC domains were also determined using the Ncoils tool implemented at the waggawagga server^35^. Several other tools were employed by the server simultaneously for high-confidence prediction. The CC domains shown in Figures 2A, 3A, 7A, and 7B were based on predictions from the Ncoils tool with a minimum window length of 21 aa.

##### Tudor domains

Tudor domains were obtained from UniProt and are listed in Table S1. In HeLa cells, 79 genes encode proteins with at least one Tudor domain, resulting in an expected frequency of Tudor domains of 0.0088.

##### RG/RGG domains

These domains were obtained from Thandapani et al., (2013)^29^ and contain at least two neighboring RG repeats or two neighboring RGG repeats. They are listed in Table S1. Among the proteins expressed in HeLa cells, 600 contain RG/RGG domains, resulting in an expected frequency of RG/RGG domains of 0.067.

##### Protein domain enrichment

To determine whether a protein domain is considered enriched among the FXR1 network-dependent interactors, we calculated the observed over expected frequency of CC, Tudor, or RG/RGG domains. The expected frequency is the frequency of domains observed in HeLa cells. The observed frequency of protein domains was obtained from the top 20% of most FXR1 network-dependent protein interactors. These proteins have the lowest log2 FC of FXR1-CC2mut/FXR1-WT. A Chi-square test was performed to test if the enrichment is statistically significant (Table S3).

### Protein sequence conservation analysis

Sequence conservation was calculated by computing the global alignment across 375 (metazoa) orthologous FXR1 sequences identified using the EggNog server^78^. Alignment was performed using Clustal Omega, and conservation was determined using the default analysis for conservation in JalView^79^.

### Gene ontology analysis

Gene ontology analysis was performed with FXR1 network dependent mRNA targets using DAVID^46^.

### mRNA abundance of FXR family proteins across cell types

The values were obtained from the Human Cell Landscape (https://db.cngb.org/HCL/data/HCL_102_average_expression.xlsx)^23^.

### TMT proteomics data analysis

For quantitative analysis, raw data files were processed using Proteome Discoverer (PD) version 2.4.1.15 (Thermo Scientific). For each of the TMT experiments, raw files from all fractions were merged and searched with the SEQUEST HT search engine with a Homo sapiens UniProt protein database downloaded on 2019/01/09 (176,945 entries). Cysteine carbamidomethylation was specified as fixed modifications, while oxidation (M), acetylation of the protein N-terminus, TMTpro (K) and TMTpro (N-term), deamidation (NQ), and phosphorylation (S, T, Y) were set as variable modifications. The precursor and fragment mass tolerances were 10 ppm and 0.6 Da, respectively. A maximum of two trypsin missed cleavages was permitted. Searches used a reversed sequence decoy strategy to control peptide false discovery rate (FDR) and 1% FDR was set as the threshold for identification.

The TMT experiment result was plotted as a volcano plot with biological significance defined as log2 fold change below −1.5 or over 1.5 and −log10 (*P* value) > 3.

### SILAC mass spectrometry data analysis

SILAC mass spectrometry data were processed using the MaxQuant software (Max Planck Institute of Biochemistry; v.1.5.3.30). The default values were used for the first search tolerance and main search tolerance—20 ppm and 6 ppm, respectively. Labels were set to Arg10 and Lys6. MaxQuant was set up to search the reference human proteome database downloaded from UniProt on January 9^th^, 2020. MaxQuant performed the search assuming trypsin digestion with up to two missed cleavages. Peptide, site and protein FDR were all set to 1% with a minimum of one peptide needed for identification but two peptides needed to calculate a protein level ratio. Ratio values of FXR1-CC2mut (H)/FXR1-WT (L) were log2-transformed (Table S3).

### iCLIP data analysis

The sequencing reads were mapped to hg38, and the number of unique CLIP reads that aligned to 5′UTRs, coding sequences (CDS), or 3′UTRs were counted. The sum of unique CLIP reads that were assigned to each specific mRNA correspond to the number of FXR1 binding sites in said mRNA. According to RNA-seq, in HeLa cells, 8901 genes are expressed with TPM values greater than 3. Their TPM values are listed in Table S1. Out of 8901 expressed mRNAs, in the iCLIP sample obtained using WT FXR1, we detected 6697 mRNAs with at least one FXR1 binding site. Among those, the top third of genes had seven or more FXR1 binding sites per mRNA and these mRNAs were considered FXR1 targets (*N* = 2327, Table S1). The total number of FXR1 binding sites in the WT sample was 66567, whereas it was 48417 in the CCmut2 sample. This supports our observation obtained from the oligo(dT) pulldown experiment that FXR1 dimerization promotes RNA binding. Among the FXR1 targets, we considered an mRNA to be network-dependent (*N* = 1223), if the number of FXR1 binding sites per mRNA decreased by at least two-fold, when comparing the WT and CC2mut samples (Fig. S5F). The remaining FXR1 targets (*N* = 1104) are considered network-independent (Table S1).

#### Correlation of mRNA features with FXR1 mRNA targets

mRNA length, CDS length and the percentage of AU (AU-content) were determined using transcripts from the Matched Annotation from the NCBI and EMBL-EBI (MANE)^80^ human version 1.2. For each gene, the transcript with the longest mRNA length was selected. Protein length was calculated by dividing CDS length by three. 3′UTR length was obtained from Ref-seq and the longest 3′UTR isoform of each gene was used (Table S1).

### Statistics

Statistical parameters are reported in the figures and figure legends, including the definitions and exact values of *N* and experimental measures (mean ± std or boxplots depicting median, 25^th^ and 75^th^ percentile (boxes) and 5% and 95% confidence intervals (error bars). Pair-wise transcriptomic feature comparisons and FRAP sample comparisons were performed using a two-sided Mann-Whitney test. Enrichment of protein domains was performed using a Chi-square test. The Pearson *P* value is reported. When showing bar plots, one-way ANOVA was performed. Statistical tests were performed on the means of the replicates.

## Supplementary Table and Video Legends

**Table S1**. FXR1 mRNA targets identified by iCLIP in HeLa cells, related to Figure 4.

**Table S2**. Protein abundance fold changes upon FXR1 knockdown in HeLa cells determined by TMT mass spectrometry in HeLa cells, related to Figure 5.

**Table S3**. FXR1 network-dependent protein interactors determined by SILAC mass spectrometry and protein domains enriched among FXR1 interacting proteins, related to Figure 6.

**Table S4**. List of oligos and nucleic acid sequences used in this study, related to STAR Methods.

**Video S1.** A time-lapse of mGFP-FXR1 full-length protein recorded at an interval of 10 seconds, related to Figures 1 and 3. Scale bar, 1 µm.

**Video S2.** A time-lapse of mGFP-FXR1 full-length protein recorded at an interval of 2 seconds, related to Figure 1. Scale bar, 1 µm.

**Video S3.** A time-lapse of mGFP-FXR1-N2 protein recorded at an interval of 10 seconds, related to Figure 1. Scale bar, 1 µm.

**Video S4.** A time-lapse of mGFP-FXR1-N1 protein recorded at an interval of 10 seconds, related to Figures 1 and 3. Scale bar, 1 µm.

**Video S5.** A time-lapse of mGFP-FXR1-N1 protein recorded at an interval of 2 seconds, related to Figure 1. Scale bar, 1 µm.

**Video S6.** A time-lapse of mGFP-FXR1-I304N protein recorded at an interval of 2 seconds, related to Figure 1. Scale bar, 1 µm.

**Video S7.** A time-lapse of mGFP-FXR1 full-length protein distribution during FRAP, related to Figure 1. Scale bar, 1 µm.

**Video S8.** A time-lapse of mGFP-FXR1-N1 protein distribution during FRAP, related to Figure 1. Scale bar, 1 µm.

**Video S9.** A time-lapse of mGFP-FXR1-N2 protein distribution during FRAP, related to Figure 1. Scale bar, 1 µm.

**Video S10.** A time-lapse of mGFP-FXR1-I304N protein distribution during FRAP, related to Figure 3. Scale bar, 1 µm.

